# An integrated single-cell atlas of human lung across the lifespan

**DOI:** 10.64898/2026.07.23.740230

**Authors:** Liangfeng Huang, Ziliang Huang, Shuo Feng, Ke Fang, Jie Wu, Ying Ao, Lijing Huang, Linyan Hu, Jin Zhang, Hao Sun, Zhichao Miao

## Abstract

The healthy adult lung is largely quiescent but can respond to injury and replace damaged or lost cells. However, the identities and origins of regenerative cell states remain poorly understood. Here we present the Human Developmental Lung Cell Atlas (HDLCA), a curated single-cell reference atlas of the respiratory system across the lifespan, integrating 253 datasets from 225 studies, and comprising over 18 million cells from 3,198 samples and 2,460 individuals across 34 anatomical locations in both health and disease. The HDLCA defines 142 consensus lung cell types, including 13 rare and 8 previously undescribed ones. Notably, we identify a rare intermediate alveolar epithelial progenitor (Int AP) population in normal lungs, with transcriptional signatures shared by alveolar type 2 (AT2) and alveolar type 1 (AT1) cells. Trajectory inference indicates that Int AP cells arise from both AT2 and *SCGB1A1*^+^ secretory cells and give rise to AT1 cells, with Hippo-YAP/TAZ signalling governing lineage specification. Int AP cells are associated with genetic susceptibility to idiopathic pulmonary fibrosis and chronic obstructive pulmonary disease, and their dysfunction may contribute to abnormal alveolar repair and regenerative failure in both diseases. The HDLCA provides a unified atlas-level reference for mapping respiratory cellular diversity, uncovering reparative cell dynamics and informing regenerative and cell-based therapeutic strategies.

## Introduction

With the continued rise in the incidence of major chronic respiratory diseases, including lung cancer, chronic obstructive pulmonary disease (COPD), and interstitial lung diseases (ILDs)^1,2^, it has become a significant global public health challenge^3^. Many chronic respiratory diseases with adult onset have origins in the lung development stage and childhood^4–6^. Reactivation of developmental programs or abnormal repair occurs in injured lungs, including aberrant epithelial, macrophage and fibroblast states in idiopathic pulmonary fibrosis (IPF) and COPD^7–13^. The healthy adult lung is largely quiescent, but stem and progenitor cell populations can be activated or re-enter the cell cycle after injury or disease to replace damaged or lost cells^14,15^. Recently, various stem and progenitor cells in both adult and developing lungs have been shown to critically contribute to lung regeneration and repair following injury, informing the development of regenerative therapies^16^. Thus, a comprehensive understanding of cellular states in developing and adult lungs across health and disease, together with the identification of reparative cell populations involved in lung regeneration, may accelerate the development of molecular, genetic and cell-based therapies for pulmonary disease.

Single-cell RNA sequencing (scRNA-seq) technology has become a powerful approach for dissecting cellular heterogeneity^17^. To date, more than 200 scRNA-seq publications on the human lung have focused largely on specific developmental stage^18–20^, anatomical regions^21,22^, or disease types^23–28^. However, individual studies remain constrained by restricted sampling locations, specific developmental time points, and distinct disease status, hindering the discovery of rare and transitional cells in health and disease. These publicly available datasets have provided invaluable resources for international consortia including the Human Lung Cell Atlas (HLCA)^29^ and LungMAP^30^, facilitating the generation of integrated atlases that advance the nexus of lung biology, technology, and medicine. These adult lung atlases present challenges in explaining the developmental origins of cellular states in health and disease, while fetal lung atlases fail to identify which developmental programs or molecular pathways are reactivated during injury and regeneration. A unified reference atlas covering the whole human respiratory system across the lifespan in health and disease is therefore needed to reveal the origins of reparative cells involved in lung regeneration.

The growing number of published scRNA-seq studies of the human lung^17–22,31,32^ together with the development of advanced computational tools^33–35^ have enabled increasingly comprehensive characterization of lung cellular composition and the identification of previously unrecognized cell types^22,24^. Whether these transitional cell populations represent bona fide regenerative intermediates or maladaptive cell states associated with failed repair remains a matter of active debate^24,32,36–41^. Recent studies have identified a cell population with regenerative potential that co-expresses markers of alveolar type 1 (AT1), alveolar type 2 (AT2) and airway secretory cells^24,32,36^. However, several evidence indicates that these cells only represent aberrant differentiation intermediates arising from dysfunctional alveolar stem cell repair responses^8,37–39,42^. These potential regenerative cell populations have been implicated in diseases such as pulmonary fibrosis^40^, COPD^31^, and lung cancer^43^. Yet, we still lack a clear understanding of the lineage specification among these potential regenerative cell populations and how they function in both health and disease, hindering the development of future regenerative therapeutic approaches for lung disease.

To address these challenges, we present here the Human Developmental Lung Cell Atlas (HDLCA), an integrated single-cell atlas of human lung across the lifespan. The HDLCA incorporates transcriptomic data from 225 publications and 2,460 individuals, covering more than 18 million cells from 34 anatomical locations across health and disease conditions, and defines 142 consensus cell types. Using this resource, we identify intermediate alveolar progenitor (Int AP) cells, a rare population in the normal adult lungs with transcriptional features shared by AT2 and AT1 cells. Int AP cells contribute to AT1 cell generation and may arise from both AT2 and *SCGB1A1*^+^ secretory cells. Int AP cells are linked to genetic susceptibility to both COPD and IPF, and exhibit disease-associated cellular states, abnormal differentiation and impaired regenerative potential in diseased lungs. Together, this large scale atlas facilitates an unbiased investigation of respiratory cell types, with implications for regenerative medicine and future cell-based therapeutic strategies for lung disease.

## Results

### Overview of the HDLCA

To build the comprehensive human developmental lung cell atlas (HDLCA), we curated, integrated and harmonized sc/snRNA-seq data from 253 publicly available datasets, including 3,198 samples from 225 publications across the lifespan and 34 distinct anatomic sites in health and disease (**Supplementary Fig. 1a**), yielding more than 18 million cells or nuclei after quality control (**Fig. 1a**-**c**). These samples include fetal (n = 181), infancy (n = 39), adolescence (n = 90), and adult stages (n = 1,928) (**Extended data Fig. 1a**). As a resource for the human respiratory system, the HDLCA is available through a web portal (dlca.gznl.org) that supports interactive data visualization and exploration. Across health and disease, most cells were derived from adults (61%), whereas fetal-derived cells comprised samples from the first trimester (29.5%; conception to 12 weeks), second trimester (52%; 13-27 weeks) and third trimester (18.5%; 28-40 weeks) (**Extended data Fig. 1b-c**).

**Fig. 1.**
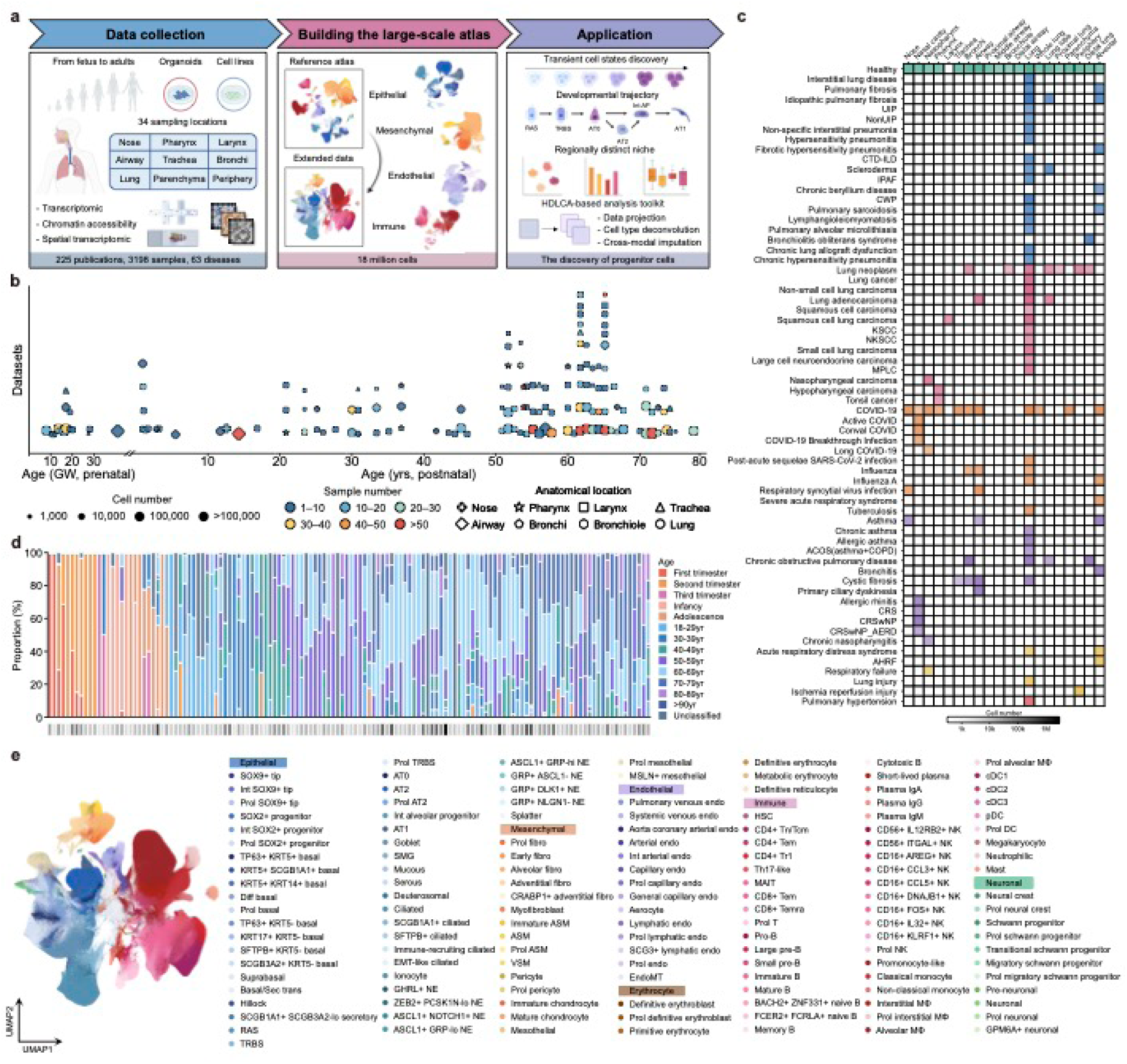
Overview of the integrated HDLCA. **a.** Schematic overview of the study design. **b.** Scatter plot showing the distribution of datasets by age in the HDLCA. Point size represents the number of cells, color indicates sample number, and shapes denote anatomical locations. **c.** Heatmap showing the distribution of disease categories across anatomical regions in the HDLCA. Color indicates disease category, and color intensity reflects the number of cells corresponding to each disease category within each anatomical region, with darker shading indicating higher cell numbers. **d.** Stacked bar plot showing age composition across studies in the HDLCA. Each bar represents one study, with colors indicating developmental stages. The bottom annotation bar indicates the number of samples per study, with darker shades corresponding to higher sample numbers. **e.** UMAP visualization of all 142 cell types in the HDLCA, with cells coloured by annotated identities.

**Extended data Fig. 1.**
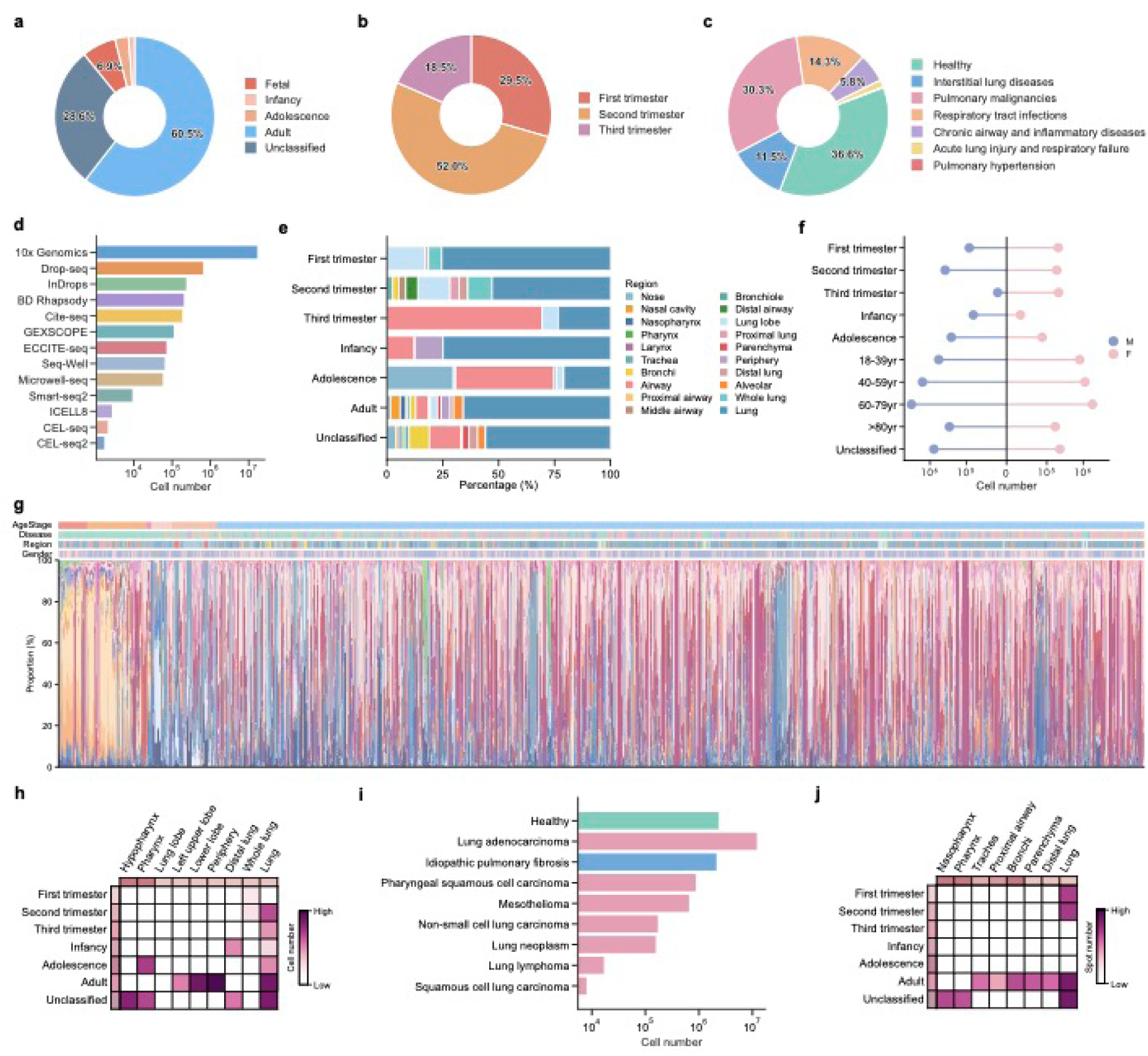
Metadata statistics of the HDLCA. **a-c**. Donut plots showing the proportions of cells across age groups (**a**), trimester stages (**b**) and disease categories (**c**) in the HDLCA. **d.** Bar plot showing the sequencing platforms included in the HDLCA. **e.** Stacked bar plot showing the proportions of anatomical regions across developmental stages. **f.** Lollipop plot showing the gender composition of samples across developmental stages by cell number. **g.** Stacked bar plot showing the proportions of cells assigned to epithelial, mesenchymal, endothelial and immune compartments in the HDLCA. **h.** Heatmap showing the sampling locations and developmental stages represented in the scATAC-seq data of the HDLCA. Color intensity denotes the number of cells corresponding to each developmental stage within individual anatomical regions, with darker shading indicating higher cell numbers. **i.** Bar plot showing the disease composition of scATAC-seq datasets included in the HDLCA. **j.** Heatmaps showing the distribution of spatial transcriptomic spots across developmental stages and anatomical regions in the HDLCA. Color intensity denotes the number of spots corresponding to each developmental stage within individual anatomical regions, with darker shading indicating higher spot numbers.

**Supplementary Fig. 1.**
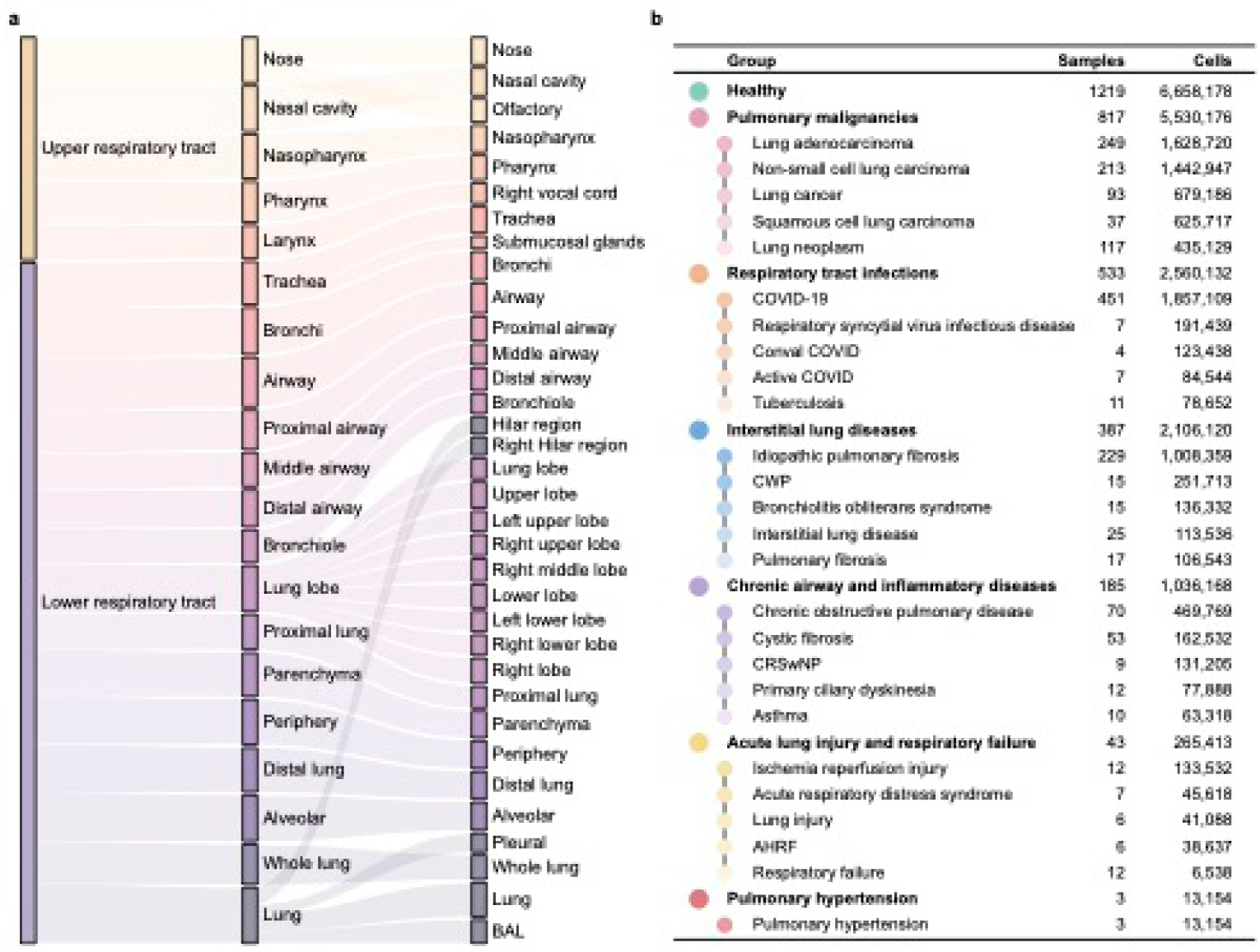
Anatomical regions and disease categories in the HDLCA. a. Sankey plot showing the hierarchical classification of anatomical regions represented in the HDLCA. b. Summary of disease categories represented in the HDLCA.

As for sampling regions, HDLCA included 34 locations such as the lung (n = 1,848), airway (n = 281), nasopharynx (n = 164), bronchi (n = 145), alveolar (n = 113), nose (n = 108), trachea (n = 96), parenchyma (n = 69), distal lung (n = 54), and nasal cavity (n = 50) (**Fig. 1b**). Transcriptomic data were generated using multiple single-cell capture systems, with most cells profiled using droplet-based platforms, including 10x Genomics, Drop-seq and inDrop. Smaller fractions were generated using microwell-based platforms (2.4%), including BD Rhapsody, GEXSCOPE, Microwell-seq, Seq-Well and ICELL8, or plate-based platforms (0.1%), including Smart-seq2, CEL-Seq and CEL-Seq2 (**Extended data Fig. 1d**). In total, cells in the HDLCA were derived from 910 healthy individuals and 1,550 diseased individuals, comprising 1,219 healthy and 1,968 diseased samples (**Supplementary Fig. 1b**). The HDLCA includes samples from female and male donors across the human lifespan, including the fetus (first, second and third trimester), infancy, adolescence and adult (**Extended data Fig. 1e-f**). A total of 63 disease types were represented in the HDLCA, among which COVID-19 was the most prevalent, followed by lung adenocarcinoma, IPF, non-small cell lung carcinoma, lung neoplasm and lung cancer (**Supplementary Fig. 1b**).

The large-scale HDLCA was constructed by extending an in-house reference atlas using the transfer learning framework scArches^44^ (see **Methods**). Automated annotation was first performed using PopV and SCimilarity algorithm (see **Methods**), followed by manual curation guided by canonical marker gene expression (**Supplementary Table 1**). The resulting annotations were aligned to the Cell Annotation Platform (CAP; https://celltype.info/). The HDLCA defines 142 consensus cell types, visualized using Uniform Manifold Approximation and Projection (UMAP) (**Fig. 1d**-**e** and **Extended data Fig. 2a**). For many cell types, biological variation across samples was largely explained by disease status and anatomical region (**Extended data Fig. 2b**). Cellular composition showed regional specificity in cell type distribution, suggesting that spatially complex cellular microenvironments are crucial for the development and regeneration of the respiratory system (**Extended data Fig. 1g**). Additionally, HDLCA incorporates spatial transcriptomic data and different modalities, such as cellular indexing of transcriptomes and epitopes by sequencing (CITE-seq) (**Extended data Fig. 1d**) and single-cell assay for transposase-accessible chromatin using sequencing (scATAC-seq) (**Extended data Fig. 1h-j**). To facilitate reference-based understanding of lung single-cell biology, we developed sc2HDLCA (**Methods**), a toolkit for annotating new datasets, imputing data across modalities, and deconvolving bulk and spatial data using the HDLCA as a reference. Hence, the HDLCA provides an integrated single-cell reference atlas and data resource at scale for the cross-conditional comparisons of respiratory cell types.

**Extended data Fig. 2.**
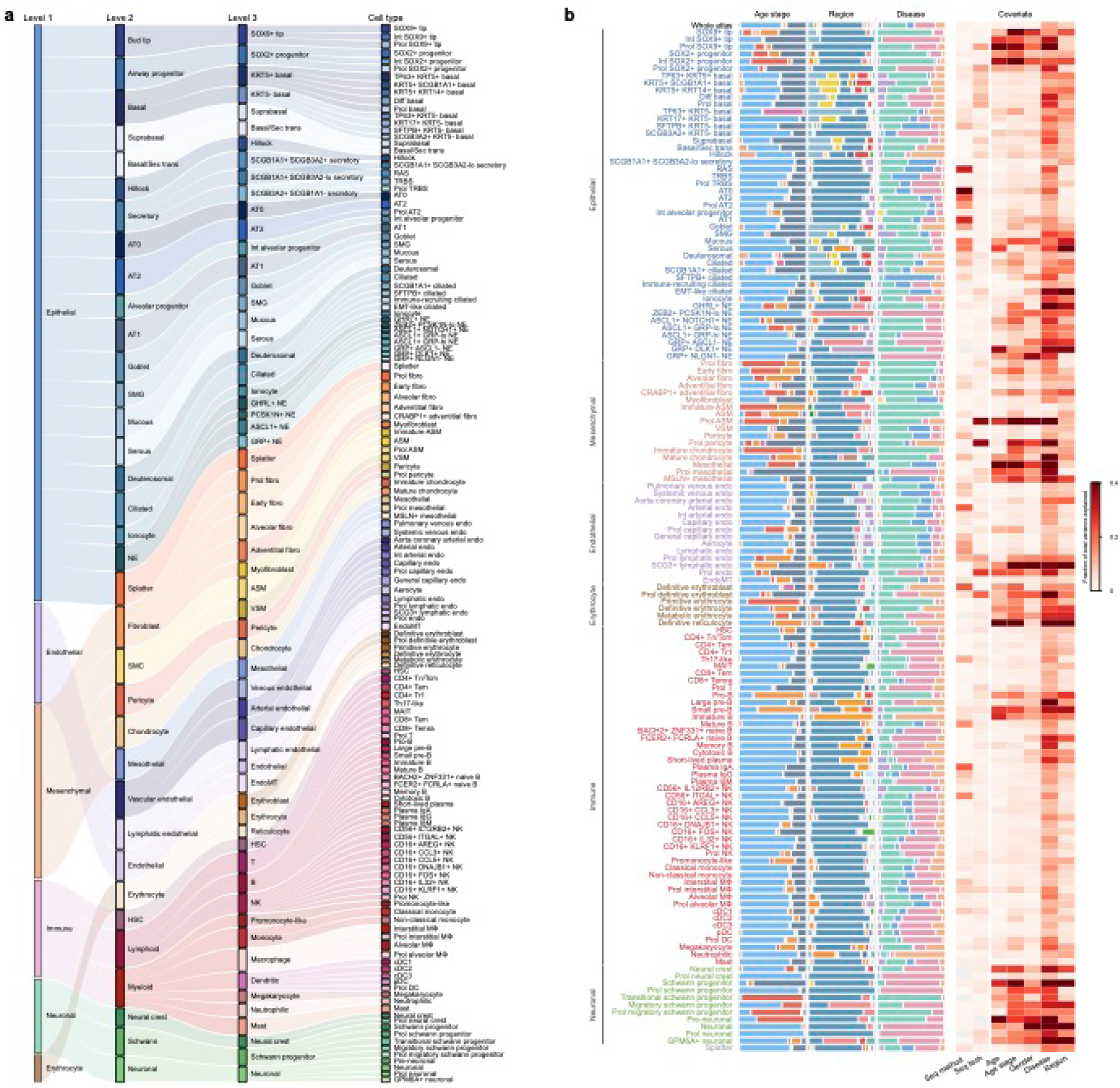
Hierarchical cell type annotation and biological variation in the HDLCA. **a.** Sankey plot showing the hierarchical cell type annotation in the HDLCA. **b.** Sample-level covariate effects across the HDLCA and annotated cell types. Rows represent the whole atlas or individual cell types grouped by major cellular compartments. Stacked bars indicate the distribution of cells across age stage, anatomical region and disease status. The heatmap shows the fraction of total inter-sample variance in the integrated embedding associated with each covariate, including sequencing method, sequencing technology, age, age stage, gender, disease status and anatomical region.

### The construction of HDLCA reference atlas

To create the HDLCA reference atlas, we integrated 13 single-cell transcriptomic datasets from 12 publications (**Supplementary Table 2**), comprising 223 normal samples from 89 donors across the human lifespan and covering 13 distinct anatomical locations from the proximal to distal lung (**Fig. 2a**). After consistent preprocessing and quality control, a total of 1,085,181 cells or nuclei were used for downstream integration and analyses (**Fig. 2b** and **Methods**). We assessed six different integration approaches (scVI^45^, scANVI^46^, scPoli^47^, Harmony^48^, SCALEX^49^ and Scanorama^50^) using single-cell integration benchmarking (scIB) framework and found that scANVI performed best for these datasets (**Extended data Fig. 3a**-**b**). To remove batch effects, scANVI was used for data semi-supervised integration based on the preliminary annotations (**Extended data Fig. 3c** and **Methods**). Integration performance was unaffected by study, donor, sample, or gender source, but developmental stage and anatomical region origin influenced integration outcomes (**Extended data Fig. 3d**-**e**). We performed clustering on the UMAP representation generated from the scANVI latent space and reannotated clusters on canonical marker gene expression (**Fig. 2c** and **Supplementary Table 1**). Evaluation using the Single-Cell Clustering Assessment Framework (SCCAF)^33^ yielded area under the curve (AUC) scores above 0.97, indicating high clustering accuracy (**Extended data Fig. 3f**).

**Fig. 2.**
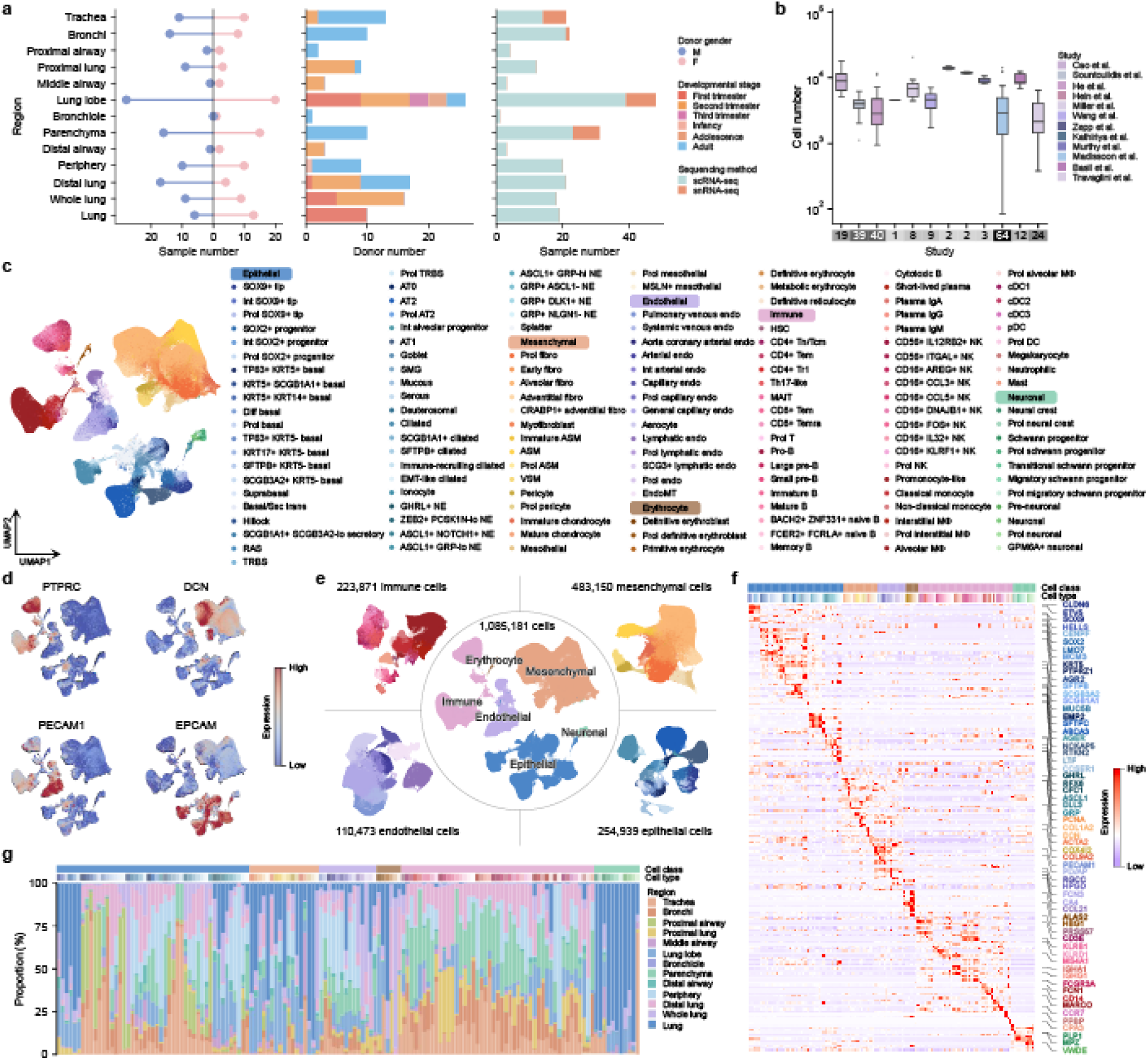
Construction of the HDLCA reference atlas. **a.** Lollipop and box plot showing the age distribution and anatomical region composition of samples used to construct the HDLCA reference atlas. **b.** Box plot showing the number of cells per sample across all studies included in the HDLCA reference atlas. The bottom annotation bar indicates the number of samples contributed by each study. Center lines indicate medians; box bounds represent the 25th and 75th percentiles; whiskers extend to values within 1.5 times the interquartile range; and points indicate outliers. **c.** UMAP representation of all 142 annotated lung cell types in the HDLCA reference atlas. **d.** UMAP visualizations showing the expression of canonical compartment-specific markers in epithelial (*EPCAM*), mesenchymal (*DCN*), endothelial (*PECAM1*) and immune (*PTPRC*) populations. Color scales indicate normalized expression levels. **e.** UMAP representation of compartment-specific cell atlases, including epithelial, mesenchymal, endothelial and immune cell populations. **f.** Heatmap showing the expression levels of representative marker genes across annotated cell populations. **g.** Stacked bar plot showing the distribution of all 142 cell types across anatomical regions.

The HDLCA reference atlas showed four transcriptionally distinct cellular compartments including the epithelial, mesenchymal, endothelial and immune lineages, as defined by the expression of compartment-specific markers *EPCAM*, *PECAM1*, *DCN* and *PTPRC*, respectively (**Fig. 2d**). To reveal the inter-compartment cellular diversity and heterogeneity, each compartment was independently re-clustered, including 254,939 epithelial, 483,150 mesenchymal, 110,473 endothelial and 223,871 immune cells (**Fig. 2e**). Based on the expression patterns of canonical marker genes, we identified 142 annotated transcriptionally distinct cell types that exhibited spatially restricted distributions across the lung, comprising 46 epithelial, 17 mesenchymal, 14 endothelial, 47 immune, 6 erythroid and 11 neuronal subsets (**Fig. 2f**-**g** and **Extended data Fig. 4a**-**d**). The HDLCA annotations were broadly consistent with the original annotations and were in agreement with the cell type classifications of HLCA and LungMAP at the coarse-grained level (**Supplementary Fig. 2a-b**).

**Extended data Fig. 3.**
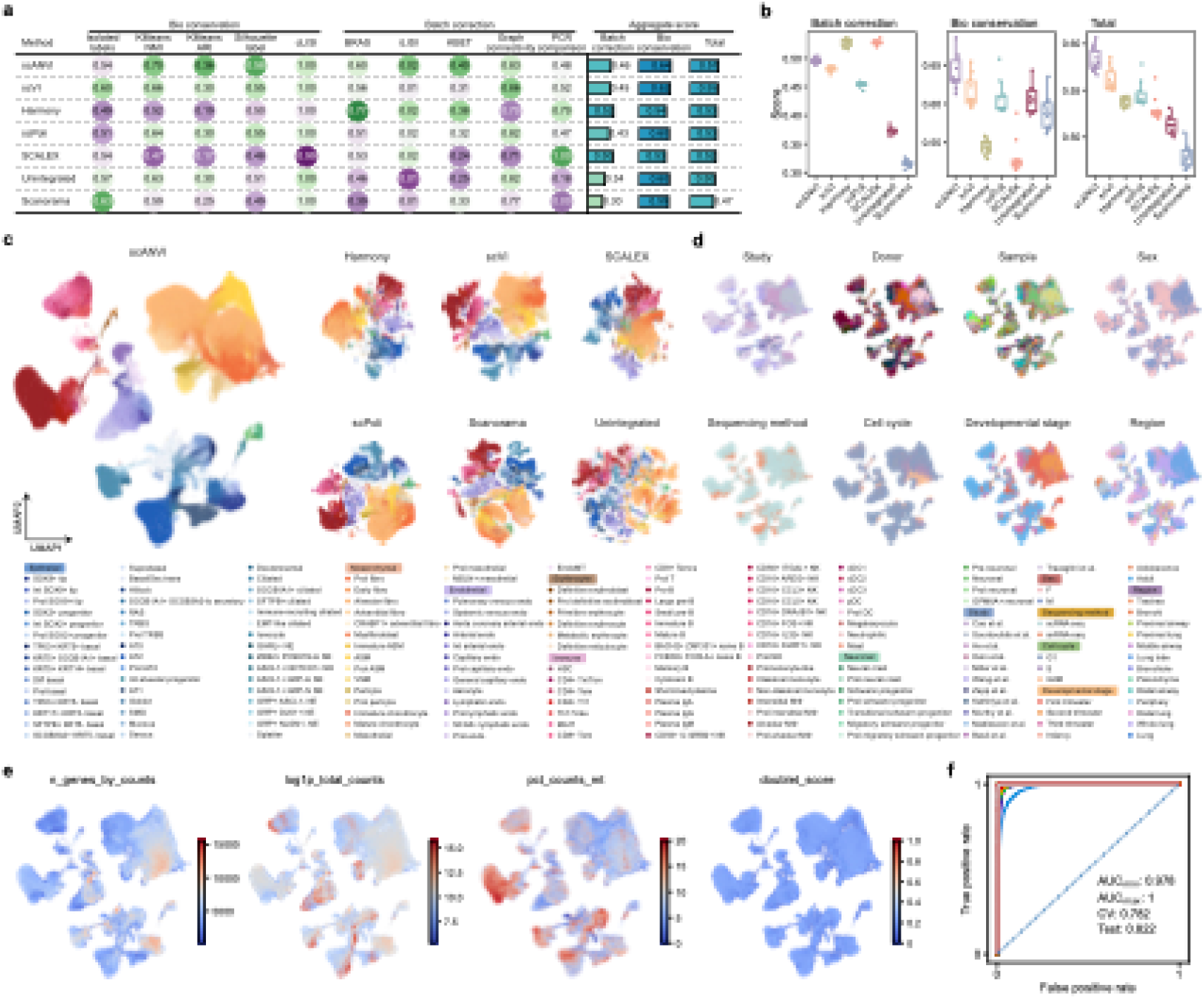
Benchmarking of data integration strategies in the HDLCA reference atlas. **a.** scIB benchmarking metrics across all evaluated data integration methods. **b.** Box plots showing biology conservation scores across integration methods. Each dot represents the benchmarking result from proportionally downsampled cells across cell types. **c.** UMAP visualization of the HDLCA dataset before integration (PCA) and after application of different data integration methods. **d.** UMAP visualization of the HDLCA reference atlas coloured by study, donor, sample, gender, sequencing platform, cell cycle, developmental stage and anatomical region. **e.** UMAP visualization showing quality-control metrics for the HDLCA reference atlas. **f.** SCCAF evaluation of clustering consistency using AUC scores.

**Extended data Fig. 4.**
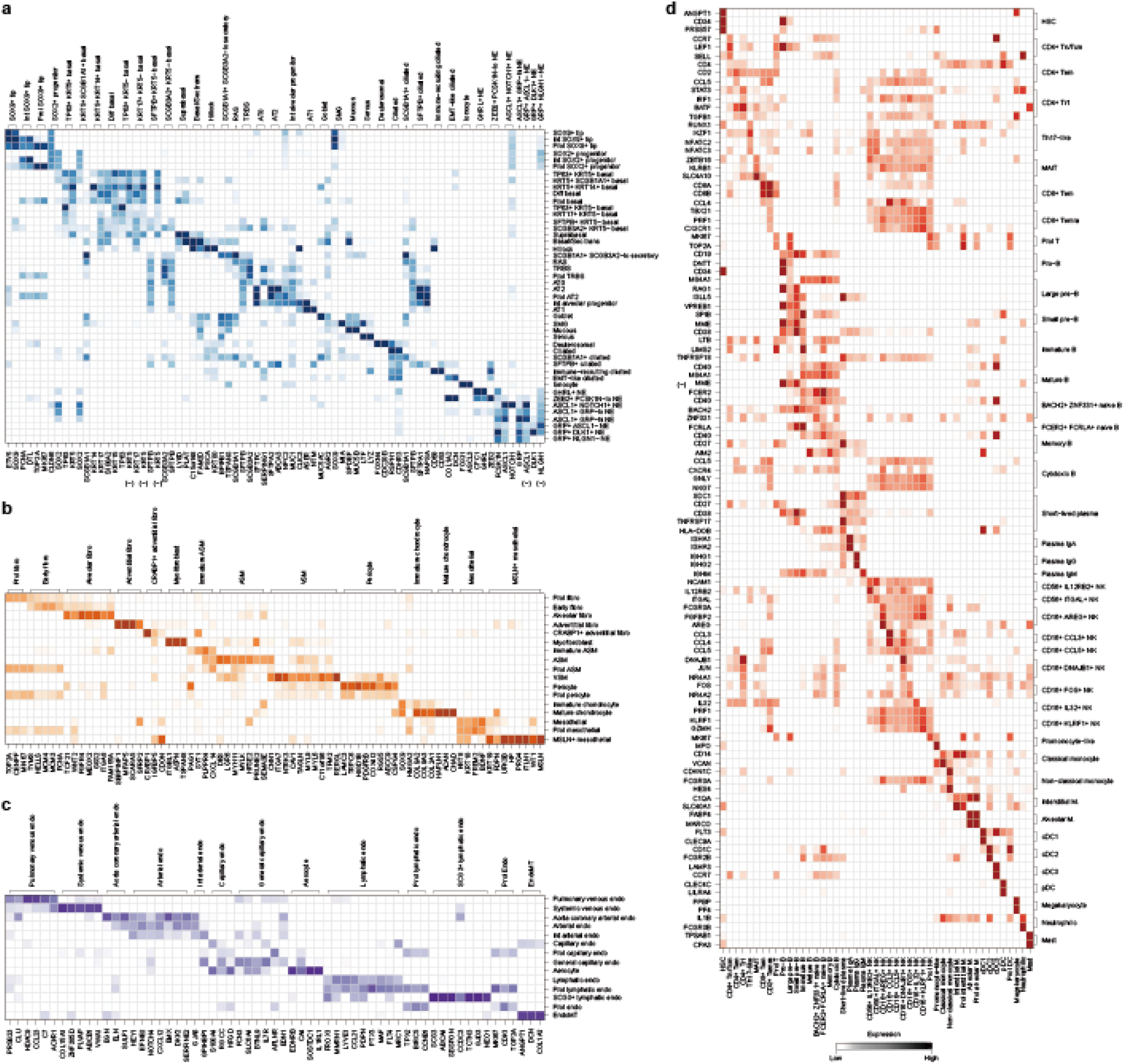
Marker gene expression for all 142 cell types in the HDLCA reference atlas. Heatmaps showing the expression of canonical marker genes across cell populations within the **a**, epithelial, **b**, mesenchymal, **c**, endothelial, and **d**, immune compartments.

**Supplementary Fig. 2.**
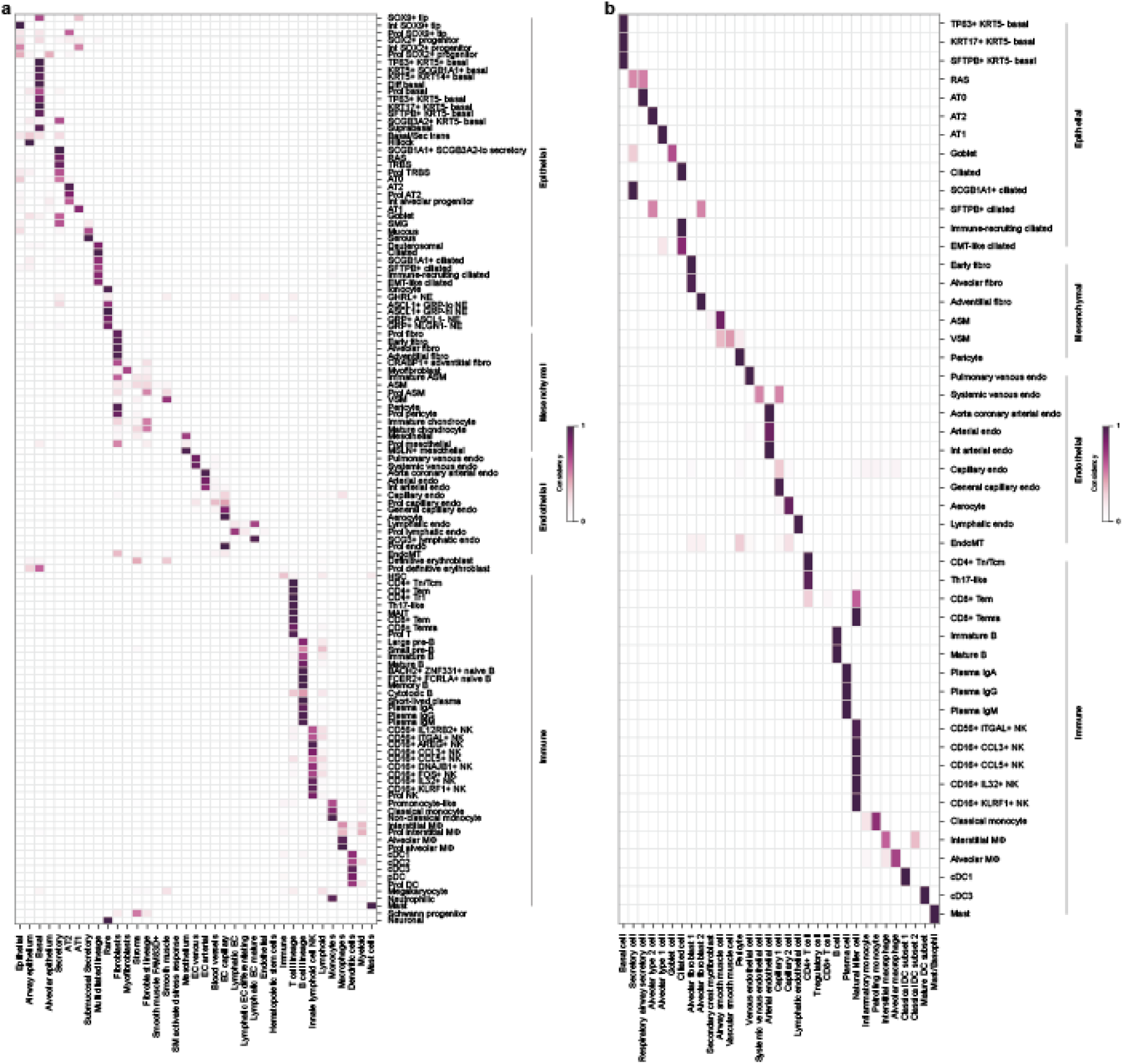
Confusion matrices assessing annotation concordance between HDLCA and existing lung cell atlases. **a.** Heatmap showing cell-type annotation concordance between HDLCA and HLCA. The x axis represents cell-type annotations from HLCA, and the y axis represents corresponding annotations from the HDLCA. **b.** Heatmap showing cell-type annotation concordance between HDLCA and LungMAP. The x axis represents cell-type annotations from LungMAP, whereas the y axis represents corresponding annotations from HDLCA.

### Identification of novel, rare and transitional cell states

The increase in total datasets from multiple sources enables the detection of rare and transitioning cell populations that might not have been found in the individual datasets^51^.

Taking advantage of the large-scale data in the HDLCA, we define the gene expression profiles of 121 canonical cell types, 13 rare and 8 previously unknown ones (**Fig. 3a**). We identified 14 proliferating cell populations in the normal lung from human adults (**Fig. 3b**). These clusters (prol *SOX2*^+^ progenitor, prol basal, prol TRBS and prol AT2) highly express common proliferation markers (*MKI67*, *TOP2A* and *BIRC5*), suggesting a low level of cellular turnover in the healthy adult lungs (**Fig. 3c** and **Extended data Fig. 5a**-**b**). We further defined ten quiescent basal cell subtypes based on the presence or absence of *KRT5* expression (**Extended data Fig. 4a**). *KRT5*-positive basal cells are mostly localized to the proximal airways, whereas *KRT5*-negative ones are largely distributed in the distal airways (**Extended data Fig. 5c**-**d**). Among these, we identified previously undescribed cell populations that co-express basal and secretory cell markers, including *KRT5*^+^/*SCGB1A1*^+^ basal, Basal/Sec trans (*KRT17*, *FAM3D* and *SCGB3A1*) and *SCGB3A2*^+^/*KRT5*^−^ basal cells (**Fig. 3d** and **Extended data Fig. 5c**). These populations may represent intermediate cellular states linking basal and secretory cell lineages. Proliferating and differentiating basal cells all derive from proximal samples, while transitioning basal cells come from both proximal and distal airways (**Extended data Fig. 5d**).

**Fig. 3.**
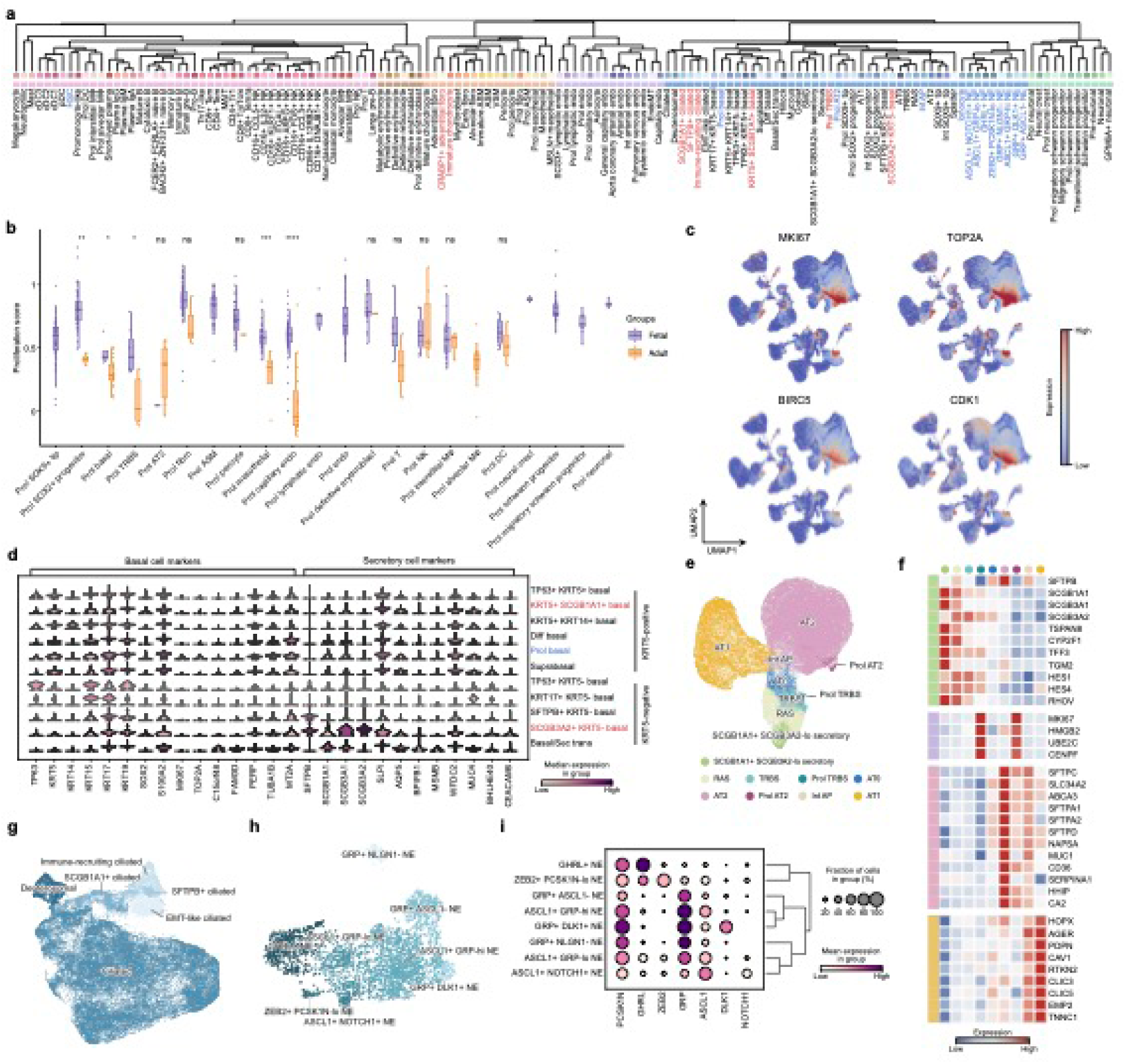
Transcriptomic profiling identifies novel, rare, and transitional cell states in the HDLCA. **a.** Hierarchical clustering dendrogram of cell populations across compartments. The upper color bar indicates annotated cell types, and the lower color bar denotes compartment identities. Font colors distinguish canonical, novel, and transitional cell states. **b.** Box plot showing proliferative gene-set scoring in proliferating cell populations derived from fetal and adult samples. Each point represents one sample. Samples with fewer than 10 cells for a given cell type were excluded. Center lines indicate medians, box limits indicate the interquartile range, and whiskers indicate 1.5 times the interquartile range. Statistical significance was calculated using a two-sided Wilcoxon rank-sum test. \**P* < 0.05, \*\**P* < 0.01, \*\*\**P* < 0.001; ns, not significant. **c.** UMAP visualization showing the expression of proliferative markers (*MKI67*, *TOP2A*, *BIRC5* and *CDK1*) in the HDLCA reference atlas. **d.** Violin plot showing the expression of marker genes across basal cell populations. Normalized expression values were used for visualization. **e.** UMAP representation of secretory and alveolar epithelial cell subsets, with cells colored according to annotated identities. **f.** Heatmap showing the expression of canonical marker genes across *SCGB1A1*^+^ *SCGB3A2*-lo secretory cells, RAS cells, TRBS cells, proliferating TRBS cells (Prol TRBS), AT0 cells, AT2 cells, proliferating AT2 cells (Prol AT2), Int AP cells and AT1 cells. **g.** UMAP representation of ciliated epithelial cell subsets coloured by annotated cell identities. **h.** UMAP representation of NE cell subsets from the HDLCA reference atlas, coloured by assigned cell identities. **i.** Dot plot showing the expression of marker genes across NE cell subsets. Dot size indicates the fraction of cells expressing each gene, and color denotes mean expression level. Normalized expression values were used for visualization.

**Extended data Fig. 5.**
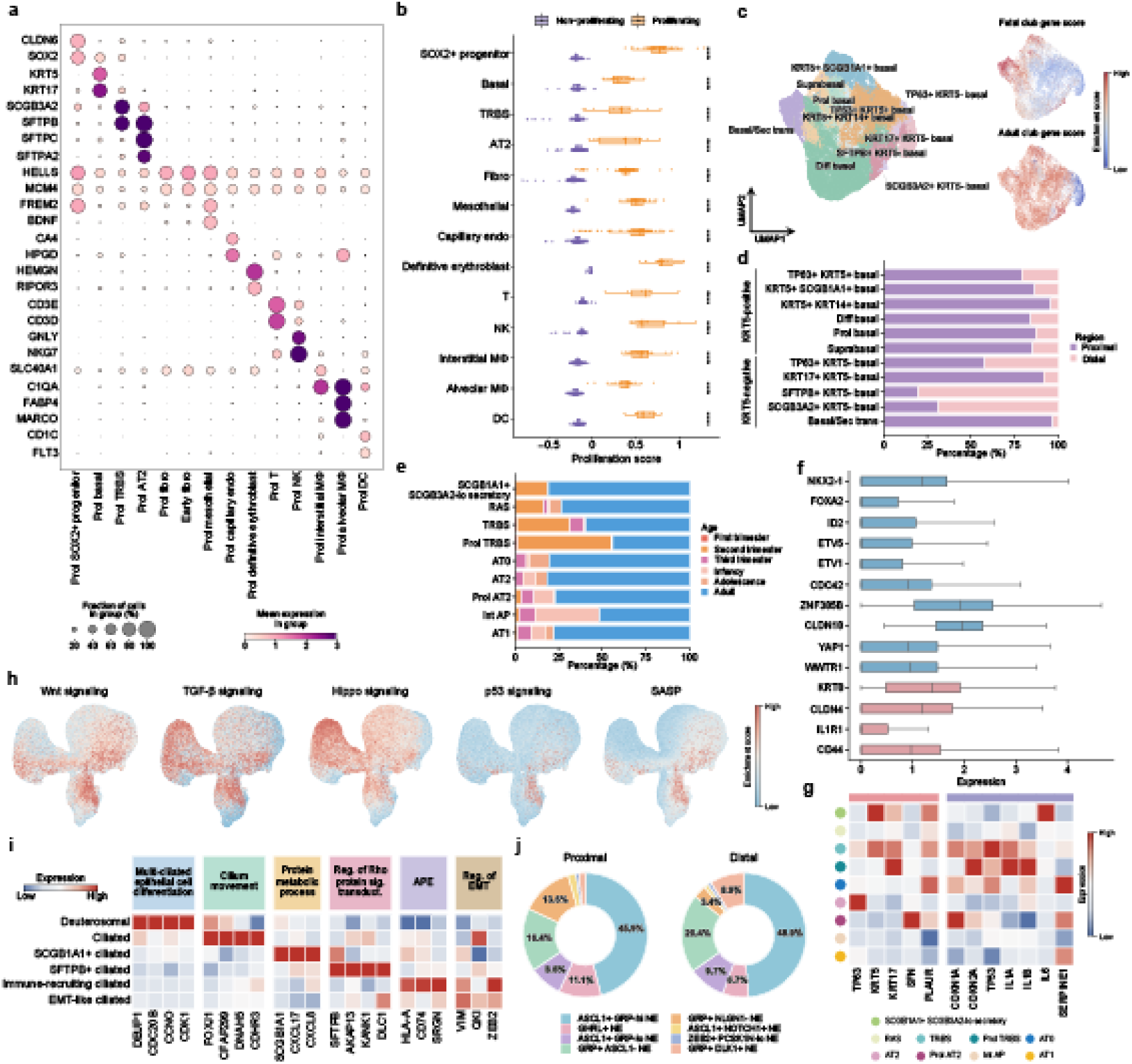
Characterization of novel, rare and transitional cell states identified in the HDLCA. **a.** Dot plot showing the expression of selected markers in 14 proliferating cell populations. Dot size indicates the fraction of cells expressing each gene, and color indicates mean normalized expression levels. **b.** Box plots showing proliferative gene set scores in proliferating and non-proliferating cell types. Each point represents one sample. Center lines indicate medians, box limits indicate the interquartile range, and whiskers indicate 1.5 times the interquartile range. Statistical significance was calculated using a two-sided Wilcoxon rank-sum test. \**P* < 0.05, \*\**P* < 0.01, \*\*\**P* < 0.001; ns, not significant. **c.** UMAP representation of basal cell subsets and secretory cell signature scores. The gene sets shown in the upper right and lower right panels were derived from the MSigDB signatures HE_LIM_SUN_FETAL_LUNG_C1_CLUB_CELL^20^ and TRAVAGLINI_LUNG_CLUB_CELL^22^, respectively. **d.** Stacked bar plot showing the spatial distribution of basal cell subtypes. **e.** Stacked bar plot showing the age composition of secretory and alveolar epithelial cell populations, including *SCGB1A1*^+^ *SCGB3A2*-lo secretory, RAS, TRBS, Prol TRBS, AT0, AT2, Prol AT2, Int AP and AT1 cells. **f.** Box plots showing the expression of selective genes in Int AP cells. Normalized expression values were used for visualization. Center lines indicate medians, box limits indicate the interquartile range, and whiskers indicate 1.5 times the interquartile range. **g.** Heatmap showing the expression of damage-associated genes and cellular senescence-associated genes across *SCGB1A1*^+^ *SCGB3A2*-lo secretory, RAS, TRBS, Prol TRBS, AT0, AT2, Prol AT2, Int AP and AT1 cells. **h.** UMAP visualization of gene set scores showing that Int AP cells have higher Wnt, TGF-β and Hippo signalling pathway scores, but lower p53 signalling and senescence-associated secretory phenotype (SASP) scores, than other distal epithelial cell types. **i.** Heatmap showing expression levels of marker genes across ciliated cell subsets. **j.** Donut plot showing the proportions of NE cell types in the proximal (left) and distal (right) airways.

We identified a rare alveolar progenitor population (Int AP; ∼0.46% of epithelial cells) in both fetal (n = 12) and adult (n = 58) normal human lungs from 31 individuals (**Extended data Fig. 5e**), which was transcriptionally distinct from previously described epithelial cell types in the HDLCA reference atlas (**Fig. 2c** and **Fig. 3e**). Int AP cells co-expressed canonical AT2 markers (*SFTPC*, *NAPSA* and *SLC34A2*) and AT1 markers (*AGER*, *CAV1* and *RTKN2*) (**Fig. 3f**). They also expressed lung developmental regulators (*NKX2-1*, *FOXA2*, *ETV1*, *ETV5* and *ID2*), the Wnt inhibitor *WIF1* and Hippo pathway terminal transcriptional activators YAP/TAZ (*YAP1* and *WWTR1*), suggesting regenerative potential (**Extended data Fig. 5f**-**h**). Compared with previously reported transitional cell states that emerge after lung injury^8,37–39,42^, Int AP cells expressed *KRT8* and *CLDN4* but lacked expression of canonical basal cell markers (*TP63*, *KRT5* and *KRT17*), stratifin (*SFN*), cell-cycle arrest genes (*CDKN1A* and *CDKN2A*) and senescence-associated markers (*TP53*, *IL1A* and *SERPINE1*) (**Extended data Fig. 5g**). Together, these findings distinguish Int AP cells from the damage-associated transitional states previously described in diseased lungs^8,37–39,42^ and support its existence as a rare cell population in the human normal lungs.

The re-clustering of ciliated cells identified six distinct clusters along the proximal-distal axis (**Fig. 3g**). We uncovered previously undescribed ciliated cell states from both developing and adult lung, including *SCGB1A1*⁺, immune-recruiting and a rare subtype termed *SFTPB*⁺ ciliated. Based on the expression of cilia and flagella-associated protein *CFAP299*, *SCGB1A1*^+^ ciliated cells express chemokines *CXCL17* and *CXCL8*, while immune-recruiting ciliated cells express MHC Class I (*HLA-A* and *HLA-B*), implying potential immune cell recruitment functions. *SFTPB*⁺ ciliated cells expressed the motile cilia transcription factor (TF) *FOXJ1* and the distal marker *SFTPB*, occupying approximately 0.09% of the cells in the HDLCA reference atlas (**Extended data Fig. 5i**). We annotated eight rare neuroendocrine (NE) cell subtypes along the airway proximal-distal axis based on the expression patterns of *ASCL1*, *GRP*, *GHRL* and *PCSK1N* (**Fig. 3h** and **Extended data Fig. 5j**). Two previously undescribed populations were identified as *CRABP1*+ adventitial fibro (*CRABP1*, *IGFBP5* and *CD248*) and immature chondrocyte (*SOX9*, *COL9A3* and *COL2A1*) in the mesenchymal compartment (**Fig. 3a** and **Extended data Fig. 4b**).

### Association of Int AP cells with pulmonary disease risk

Given that treatment for heterogeneous pulmonary diseases such as COPD and IPF remains largely homogeneous, resulting in significant variations in clinical outcomes, precision-targeted therapy promises to improve this situation^52^. To identify disease-associated cell states in the HDLCA, we applied gsMap^53^ to integrate genome-wide association study (GWAS) summary statistics for 17 pulmonary diseases with 68 10x Genomics Visium spatial transcriptomic datasets, revealing a significant association of Int AP cells with COPD and IPF (**Fig. 4a** and **Supplementary Table 3**). Genes associated with COPD-susceptibility including *SFTPB*^54^, *NPNT*, *SERPINA1* and *HHIP*^55^, as well as genes linked to IPF^56^ (*SFTPC*, *SFTPA2*, *ABCA3* and *ATP11A*), predominantly expressed in the distal epithelial cell populations (**Fig. 4b**).

**Fig. 4.**
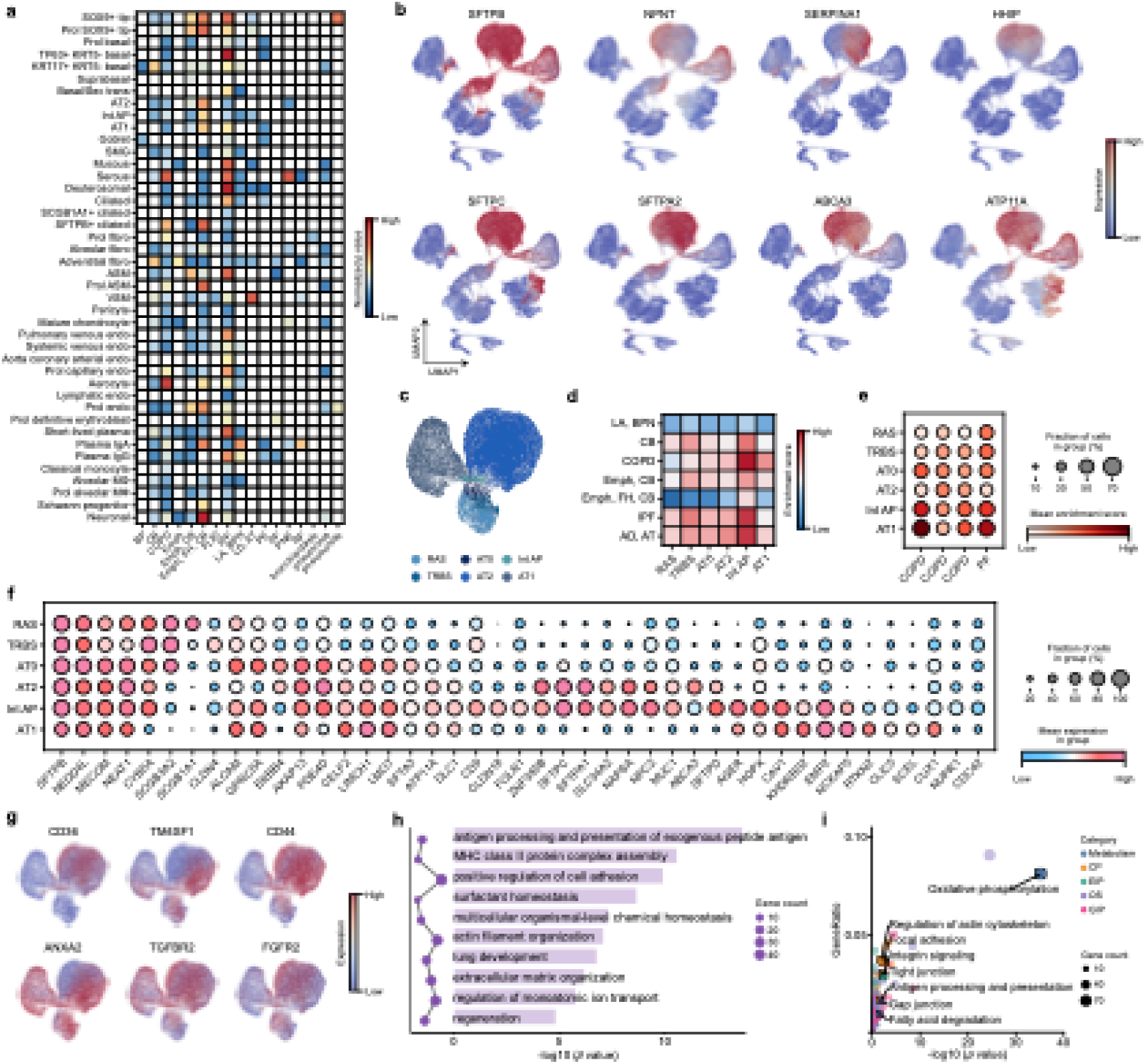
Int AP cells exhibit genetic susceptibility to both COPD and IPF. **a.** Heatmap showing aggregated *P* values across pulmonary diseases. Aggregated *P* values were calculated using the Cauchy combination test to aggregate *P* values from individual spots within each cell type. Statistical significance was calculated using a one-sided Z-test. Abbreviations: BP, bacterial pneumonia; CB, chronic bronchitis; COPD, chronic obstructive pulmonary disease; Emph, emphysema; Emph, CB, emphysema with chronic bronchitis; Emph, FH, CB, emphysema with family history and chronic bronchitis; FLID, fungal lung infectious disease with pneumonia; IPF, idiopathic pulmonary fibrosis; LA, BPN, lung abscess and bronchopneumonia; LD, AT, lung disease with abnormal thrombosis; PE, pulmonary edema; PF, pulmonary fibrosis; PNE, pneumoconiosis; RF, respiratory failure. **b.** UMAP visualization showing expression patterns of representative susceptibility genes associated with COPD and IPF. **c.** UMAP representation of distal epithelial cell populations, with cells colored according to annotated identities. **d.** Heatmap demonstrating gsMap-identified risk gene set scores across seven pulmonary diseases. **e.** Dot plot showing the association of Int AP cells with COPD and IPF based on scDRS^57^ analysis. COPD GWAS summary statistics were downloaded from the GWAS Catalog, with accession IDs GCST90044074, GCST90436220 and GCST90468119 shown from left to right. **f.** Dot plot showing the expression of canonical marker genes across RAS, TRBS, AT0, AT2, Int AP and AT1 cell populations. **g.** UMAP visualization showing the expression of Int AP signature genes, including *CD36*, *TM4SF1*, *CD44*, *ANXA2*, *TGFBR2* and *FGFR2*. **h.** Bar plot showing GO enrichment of DEGs in Int AP cells identified by Wilcoxon rank sum test. Statistical significance was assessed using a one-sided hypergeometric test, with *P* values adjusted using the Benjamini–Hochberg method. **i.** Bubble plot showing pathway enrichment analysis of Int AP DEGs identified by Wilcoxon rank sum test. Each bubble represents one pathway, with bubble size indicating the number of enriched genes. Statistical significance was assessed using a one-sided hypergeometric test, with *P* values adjusted using the Benjamini–Hochberg method.

These cells express distal lung markers (*SFTPB*, *MECOM* and *NEDD4L*) were re-clustered into distinct RAS (*SCGB1A1*^+^/*SCGB3A2*^+^), TRBS (*SCGB3A2*^+^/*SCGB1A1*^−^/*SFTPC*^−^), AT0 (*SCGB3A2*^+^/*SFTPC*^+^), AT2 (*SFTPC*^+^/*SLC34A2*^+^), Int AP (*SFTPC*^+^/*AGER*^+^) and AT1 (*AGER*^+^/*RTKN2*^+^) populations (**Fig. 4c** and **Supplementary Fig. 3a-c**). GWAS-identified genetic susceptibility gene set scoring and single-cell disease-relevance score (scDRS)^57^ analysis further validate the significant contribution of Int AP cells to disease susceptibility in both COPD and IPF (**Fig. 4d-e**). We next examined the expression of disease risk variant-associated genes, including *FAM13A*, *SFTPD*, *AGER*, *PID1*, *AKAP13*, *MUC1* and *HLA-A*, which were highly expressed in Int AP cells (**Extended data Fig. 6a-b**). Twenty-six genes encoding viral receptors were highly expressed in at least one distal epithelial cell type. Notably, Int AP cells exhibited elevated expression of genes implicated in viral susceptibility, including human rhinovirus receptors (*ICAM1* and *LDLR*), the coxsackievirus and adenovirus receptor *CXADR*, the SARS-CoV-2 entry-associated integrin *ITGB6* and additional viral interaction factors (*CD151*, *CD46* and *CD55*) (**Extended data Fig. 6c**).

**Extended data Fig. 6.**
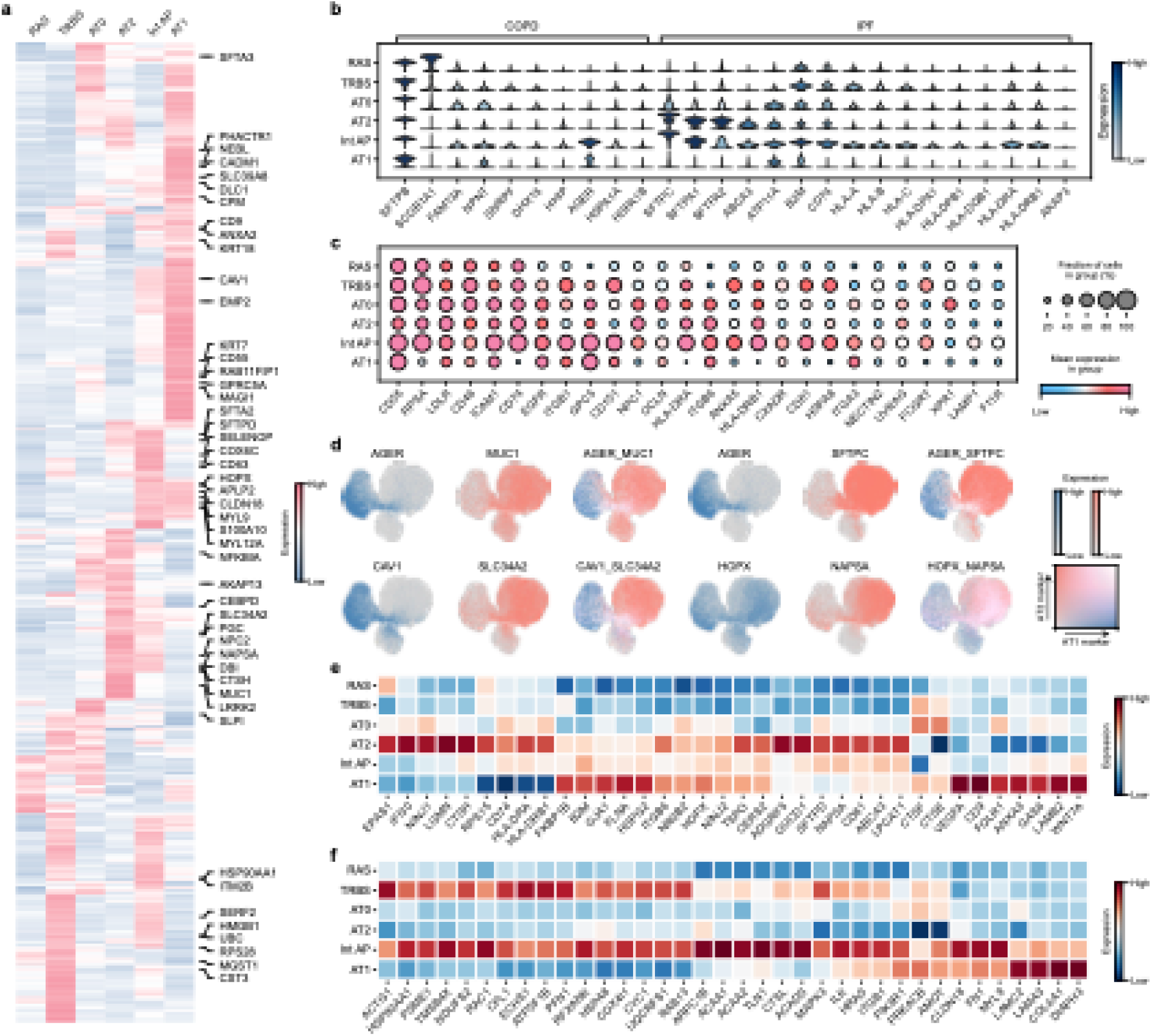
Transcriptomic signature of Int AP cells. **a.** Heatmap showing the expression of COPD- and IPF-associated risk variant genes identified by gsMap^53^ algorithm. **b.** Violin plot showing the expression of representative genes located within COPD- and IPF-susceptibility loci. **c.** Dot plot showing average expression of representative viral receptors across distal epithelial cell populations. **d.** UMAP visualization showing co-expression of AT1 markers (*AGER*, *CAV1* and *HOPX*) and AT2 markers (*SFTPC*, *SLC34A2* and *NAPSA*) in Int AP cells. **e.** Heatmap showing the expression of genes associated with the pathways presented in Fig. 4h (**e**) and Fig. 4i (**f**) within the distal epithelial cells.

**Supplementary Fig. 3.**
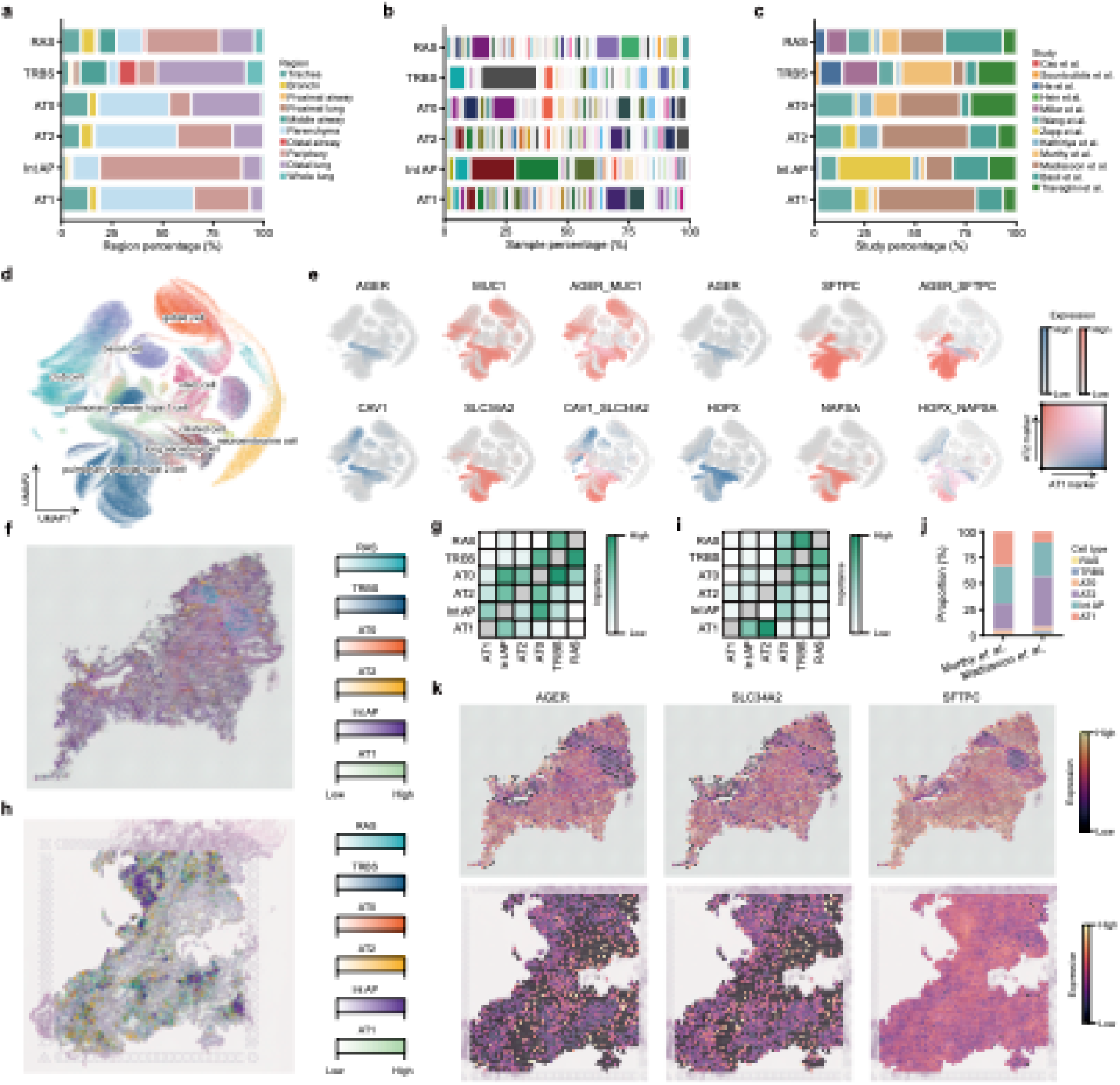
Detection of Int AP cells in organoids and spatial transcriptomics datasets. **a-c**. Stacked bar plots showing the distribution of anatomical locations (**a**), samples (**b**) and publications (**c**) across RAS, TRBS, AT0, AT2, Int AP and AT1 populations. **d.** UMAP visualization of the HLOCA^58^ with cells coloured by original cell type annotation. **e.** UMAP visualization showing co-expression of AT1 markers (*AGER*, *CAV1* and *HOPX*) and AT2 markers (*SFTPC*, *SLC34A2* and *NAPSA*) in HLOCA^58^. **f.** Cell2location mapping on a 10x Visium section of distal airway from Murthy *et al*.^59^, showing the spatial distribution of RAS, TRBS, AT0, AT2, Int AP and AT1 cell types. Spatial deconvolution was performed using the HDLCA reference atlas as the reference. **g.** Heatmap showing the median importance of cell type abundance in predicting the abundance of other cell types within a spot, as inferred by mistyR^61^ algorithm using spatial transcriptomic data from Murthy *et al*.^59^ **h.** Cell2location mapping on a 10x Visium section of distal airway from Madissoon *et al*.^60^, showing the spatial distribution of RAS, TRBS, AT0, AT2, Int AP and AT1 cell types. Spatial deconvolution was performed using the HDLCA reference atlas as the reference. **i.** Heatmap showing the median importance of cell type abundance in predicting the abundance of other cell types within a spot, as inferred by mistyR^61^ algorithm using spatial transcriptomic data from Madissoon *et al*.^60^ **j.** Stacked bar plot shows the cell proportions among distal epithelial cell types in spatial datasets from Murthy *et al*.^59^ and Madissoon *et al*.^60^ **k.** Spatial sections showing the expression of AT1 (*AGER*) and AT2 (*SLC34A2* and *SFTPC*) marker genes in spatial datasets from Murthy *et al*.^59^ and Madissoon *et al*.^60^

Int AP cells co-express AT2 and AT1 markers, together with the alveolar epithelial cell surface marker *CD36*, alveolar progenitor cell marker *TM4SF1*, stem cell signaling factor *CD44*, annexin A2 (*ANXA2*), transforming growth factor beta receptor 2 (*TGFBR2*) and fibroblast growth factor receptor 2 (*FGFR2*), suggesting its in vivo self-renewal capacity and differentiation potential into AT1 cells (**Fig. 4f-g** and **Extended data Fig. 6d**). We further validated the presence of Int AP cells in the human lung organoid cell atlas (HLOCA)^58^ (**Supplementary Fig. 3d-e**) and in spatial transcriptomic datasets generated by Murthy *et al*.^59^ (**Supplementary Fig. 3f-h**) and Madissoon *et al*.^60^ (**Supplementary Fig. 3i-k**). Gene ontology (GO) analysis showed that differentially expressed genes (DEGs) in Int AP cells were enriched in MHC-II antigen-presenting genes (*HLA-DRA* and *HLA-DRB1*),surfactant homeostasis genes (*NAPSA*, *SFTPD* and *LPCAT1*) and regeneration-associated genes (*HOPX*, *FOLR1* and *ANXA3*) (**Fig. 4h**, **Extended data Fig. 6e** and **Supplementary Table 4**). Kyoto Encyclopedia of Genes and Genomes (KEGG) pathway enrichment analysis further indicated that Int AP cells were metabolically active (oxidative phosphorylation), showed features associated with cell-matrix adhesion and migration (focal adhesion, integrin signaling and regulation of actin cytoskeleton), epithelial barrier function (tight junction and gap junction) and immune regulation (antigen processing and presentation) (**Fig. 4i, Extended data Fig. 6f** and **Supplementary Table 4**). Collectively, these findings indicate that Int AP cells combine immunological and regenerative features and are susceptible to both COPD and IPF.

### Dual-origin Int AP cells generate AT1 cells

Using lineage trajectory inference tools, including scVelo^62^, Monocle 3^63^, PAGA^64^, Diffusion Map^65^, Slingshot^66^, Palantir^67^ and SLICE^68^, we found that Int AP cells have the potential to give rise to AT1 cells and originate from both AT2 and *SCGB3A2*⁺ secretory cells (**Fig. 5a-c**). We validated the unipotent state of Int AP cells, as indicated by higher CytoTRACE2 scores (**Fig. 5d-e**). Quantification of differentiation potential using single-cell entropy (scEntropy) score revealed that Int AP cells occupy an intermediate state between AT2 and AT1 cells (**Fig. 5f** and **Extended data Fig. 7a**). In both UMAP and diffusion map visualizations, Int AP cells were positioned between AT1 and AT2 cells and adjacent to AT0 cells (**Fig. 5g**). Slingshot analysis indicated that RAS marks the starting point of differentiation for these *SFTPB*-positive distal epithelial cells, revealing two distinct lineage trajectories arise: RAS→TRBS→AT0→Int AP→AT1 (Lineage 1) and RAS→TRBS→AT0→AT2 (Lineage 2) (**Fig. 5g-h** and **Extended data Fig. 7b-c**). With increasing pseudotime, the expression levels of secretory cell marker *SCGB1A1* progressively decreased, whereas alveolar epithelial lineage markers (*SFTPC*, *ABCA3*, *AGER* and *CLIC5*) gradually increased, validating the accuracy of the putative trajectories (**Fig. 5i-j**).

**Fig. 5.**
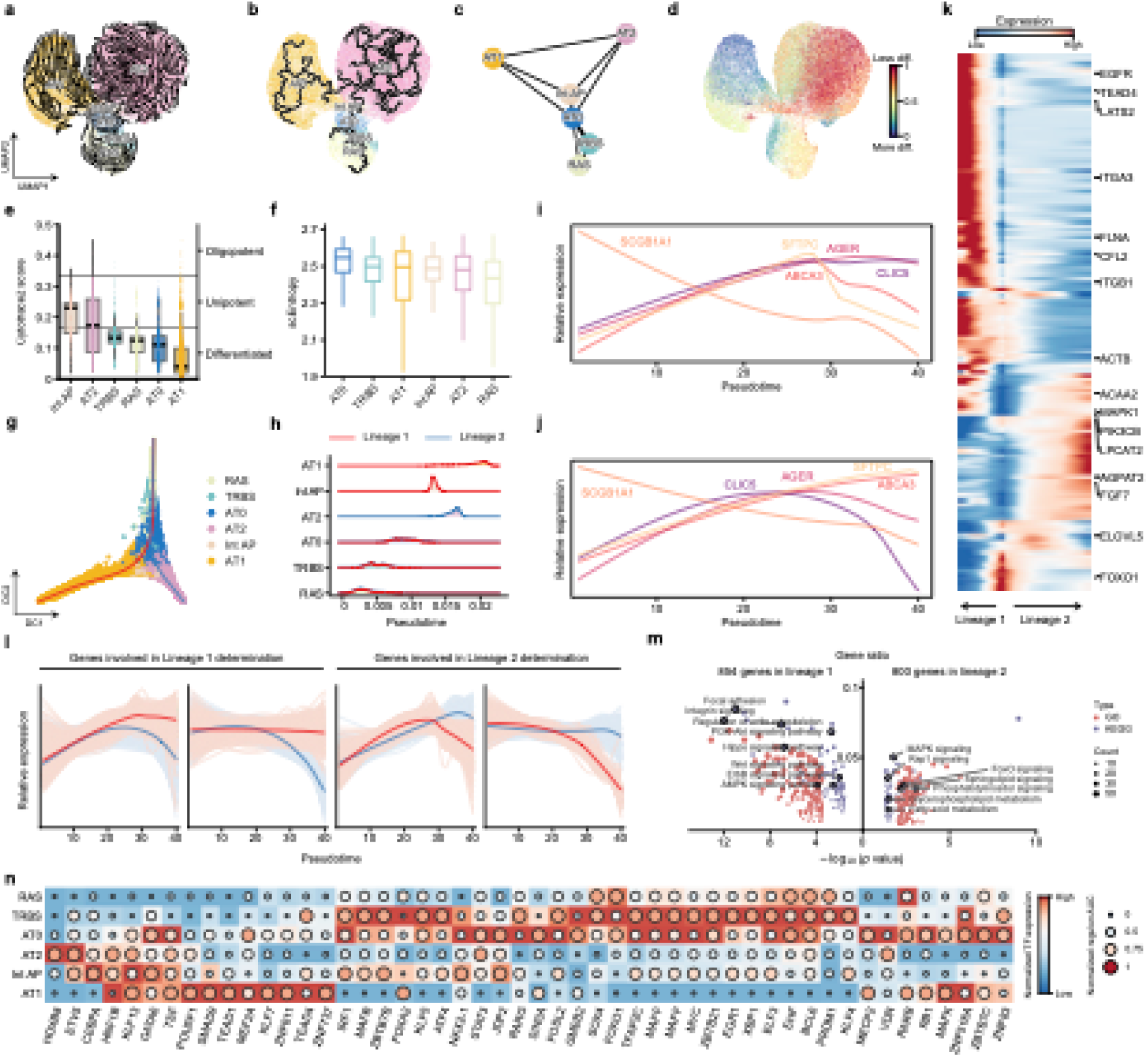
RAS- and AT2-derived Int AP cells can differentiate into AT1 cells. **a.** RNA velocity inferred by scVelo^62^ in the UMAP-based embedding space. Each point denotes a single cell and is coloured by cell identity. **b.** UMAP representation of the Monocle 3 trajectory. **c.** PAGA^64^ analysis of distal epithelial cell differentiation trajectories. Each point denotes one cell type and is coloured by annotated identity. **d.** UMAP representation of distal epithelial cells coloured by differentiation state inferred using CytoTRACE2^72^. **e-f**. Box plots showing the distribution of CytoTRACE2 scores (**e**) and scEntropy scores (**f**) across RAS, TRBS, AT0, AT2, Int AP and AT1 cell populations. **g**. Trajectory inference using Slingshot^66^ based on diffusion map embedding space. Lines indicate the predicted lineage trajectories. **h**. Ridge plot showing the distribution of pseudotime values across cell types. Line color indicates predicted lineage trajectories. **i-j**. Line plots showing expression dynamics of selected genes across 40 pseudotime-ordered bins along Lineage 1 (**i**) and Lineage 2 (**j**). The x-axis represents evenly divided pseudotime intervals and the y-axis represents normalized expression levels. **k.** Heatmap showing expression dynamics of trajectory-associated genes along pseudotime. **l.** Clustering line plot showing expression patterns of dynamic genes across 40 pseudotime-ordered points along each lineage. Lines are coloured by lineage trajectory. **m.** Bubble plot showing enriched GO biological process terms and KEGG pathways in Lineage 1 and Lineage 2. Bubble size indicates the number of enriched genes. The top 160 GO terms and top 90 KEGG pathways were ranked by *P* value for each lineage. Statistical significance was assessed using a one-sided hypergeometric test, with *P* values adjusted using the Benjamini–Hochberg method. **n.** Normalized TF expression levels and AUC scores of selected regulons across cell types.

**Extended data Fig. 7.**
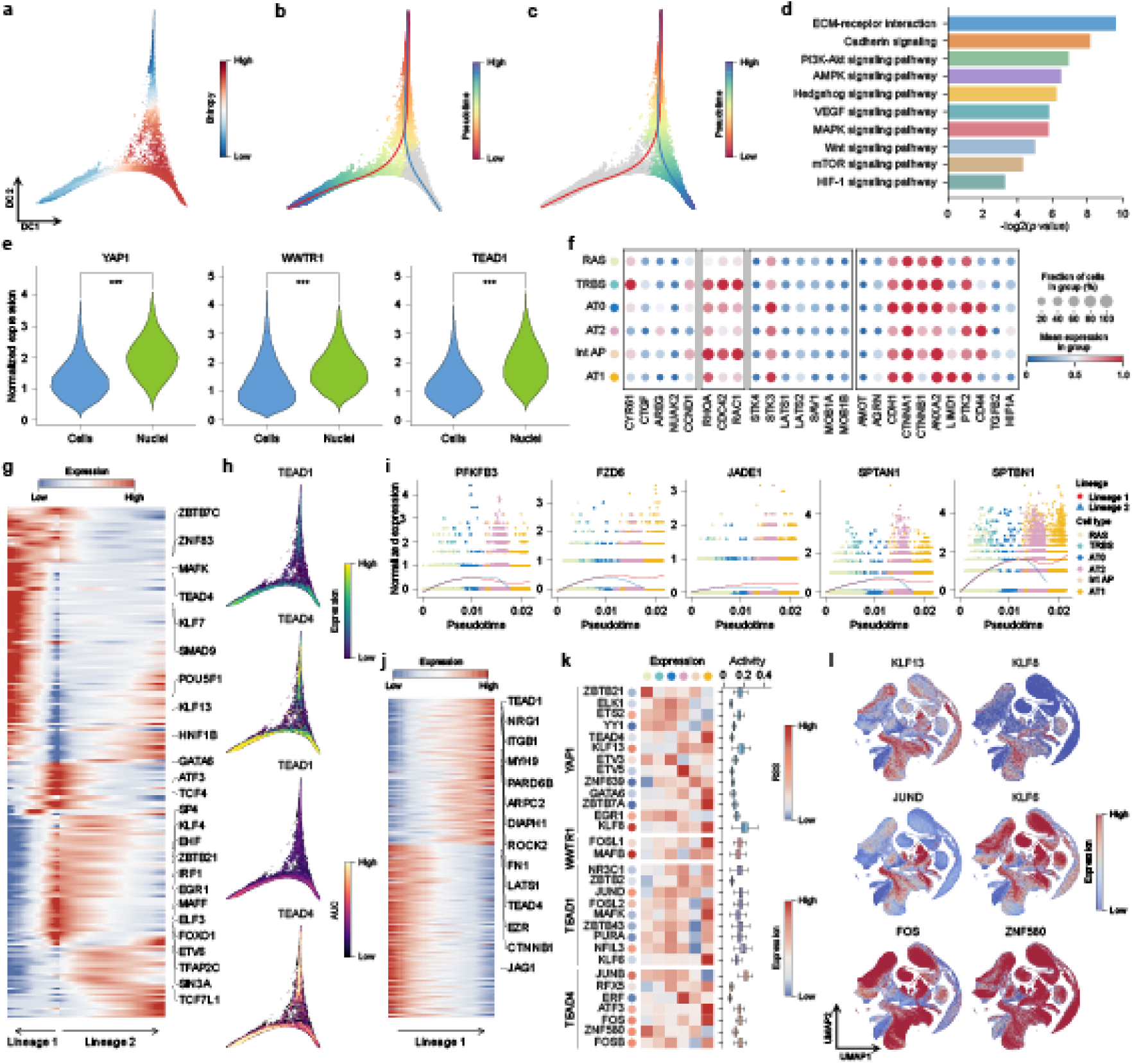
Hippo-YAP/TAZ signalling is required for Int AP lineage determination. **a-c**. Diffusion map visualization of RAS, TRBS, AT0, AT2, In AP and AT1 cells, coloured by entropy scores (**a**), and pseudotime values along Lineage 1(**b**) and Lineage 2 (**c**). Lines indicate inferred lineage trajectories. **d.** KEGG pathway enrichment analysis of genes enriched along Lineage 1. Significantly enriched pathways are ranked by *P* value. Statistical significance was assessed using a one-sided hypergeometric test, with *P* values adjusted using the Benjamini–Hochberg method. **e.** Violin plots comparing *YAP1*, *WWTR1* and *TEAD1* expression in Int AP cells from the HDLCA reference atlas between scRNA-seq and snRNA-seq datasets. **f.** Dot plot showing the expression of Hippo-YAP signaling pathway genes in the HDLCA reference atlas. **g.** Heatmap showing the dynamic expression patterns of 242 TFs inferred by SCENIC^71^ along the pseudotime differentiation trajectories. **h.** Diffusion map representation of *TEAD1* and *TEAD4* regulon expression and activities associated with trajectory determination. **i.** Scatter plots with smoothed expression curves showing the dynamic expression patterns of *TEAD1* target genes associated with trajectory determination, including *PFKFB3*, *FZD6*, *JADE1*, *SPTAN1* and *SPTBN1*. Each point represents a single cell and is coloured by cell identity. **j.** Heatmap showing dynamic expression changes of *TEAD4* target genes associated with Lineage 1 determination along pseudotime. **k.** Heatmap showing representative TF expression levels, regulon activities and regulon specificity scores (RSS). **l.** Dot plot showing the expression of Hippo-YAP signaling pathway genes in HLOCA^58^.

To elucidate the molecular pathways of RAS cell differentiation, we applied the tradeSeq^69^ algorithm and identified 1,654 genes with lineage-specific changes in expression patterns along pseudotime trajectories (**Fig. 5k-l** and **Supplementary Table 5**). Genes activated in Lineage 1 were enriched for regulation of actin cytoskeleton, focal adhesion, integrin signaling, and hippo signaling pathway, whereas genes activated in Lineage 2 were enriched for MAPK signaling, FoxO signaling and pathways associated with surfactant production and metabolic activity (**Fig. 5m**). Furthermore, we found that several pathways (Wnt, PI3K-Akt and AMPK signaling) regulating the activity of Hippo pathway effectors YAP/TAZ were involved in Lineage 1 differentiation, suggesting that crosstalk between the Hippo pathway and these pathways contributes to RAS-to-Int AP differentiation (**Extended data Fig. 7d**). We also detected YAP/TAZ (*YAP1* and *WWTR1*) expression in Int AP cells from snRNA-seq, together with upregulation of Rho GTPases (*RHOA*, *CDC42* and *RAC1*), focal adhesion kinase (*PTK2*), and YAP/TAZ-TEAD target genes (*CYR61* and *CTGF*). These findings suggest that Int AP cells may represent an additional alveolar epithelial mechanosensor beyond AT1 cells^70^ and that Hippo-YAP/TAZ signalling contributes to Int AP lineage specification (**Extended data Fig. 7e-f**).

To establish the gene regulatory networks (GRNs) governing alveolar epithelial cell lineage specification, we used SCENIC^71^ to infer regulon activity based on the HDLCA reference atlas. We identified known master regulators of lung morphogenesis (*NKX2-1* and *HNF1B*), alveolarization-essential factor *FOXA2*, the key TF for airway secretory cell differentiation into AT2 cells (*FOSL2*), AT2 cells proliferation-promoting and phenotype-maintaining factor *ETV5*, and cell-type-specific regulons including *MYC* and *KLF4* as potential lineage-determining factors (**Fig. 5n**, **Extended data Fig. 7g** and **Supplementary Table 5**). As expected, YAP/TAZ target TFs (TEAD1 and TEAD4) and downstream genes (*ROCK2*, *MYH9* and *ITGB1*) were enriched in Int AP cells, indicating a key role for hippo signalling in regulating alveolar epithelial differentiation (**Extended data Fig. 7h-j**). Putative regulons controlling YAP/TAZ-TEAD activity, including *KLF13*, *JUND* and *ZNF580*, also showed elevated RNA expression and transcriptional regulatory activity in Int AP cells (**Extended data Fig. 7k**). This observation is consistent with the expression patterns of these TFs in Int AP cells identified in the organoid datasets from HLOCA^58^(**Extended data Fig. 7l**).

### Dysfunction of Int AP cells in both COPD and IPF

Persistent injury and loss of the alveolar epithelial cells are fundamental pathological features shared by COPD and IPF^73^. To characterize disease-associated molecular and cellular dynamics in Int AP cells, we constructed an integrated diseased lung cell atlas from patients with COPD and IPF. We then identified distal epithelial cell populations, including RAS, TRBS, AT0, AT2, Int AP and AT1 cells (**Fig. 6a-b**). Within this integrated atlas, we identified 681 Int AP cells from 31 COPD samples across 9 studies and 2,060 Int AP cells from 131 IPF samples across 14 studies (**Fig. 6c** and **Extended data Fig. 8a**). Comparative transcriptomic analysis of Int AP cells from 31 control individuals and 128 patients with COPD (n = 31 donors) or IPF (n = 97 donors) revealed both shared and disease-specific molecular features. Among 2,094 DEGs identified in IPF and 1,555 DEGs identified in COPD, 997 genes showed concordant expression changes, including 390 commonly upregulated genes and 605 commonly downregulated genes (**Fig. 6d** and **Supplementary Table 6**). These findings suggest that IPF and COPD share transcriptional alterations associated with epithelial injury and impaired alveolar repair.

**Fig. 6.**
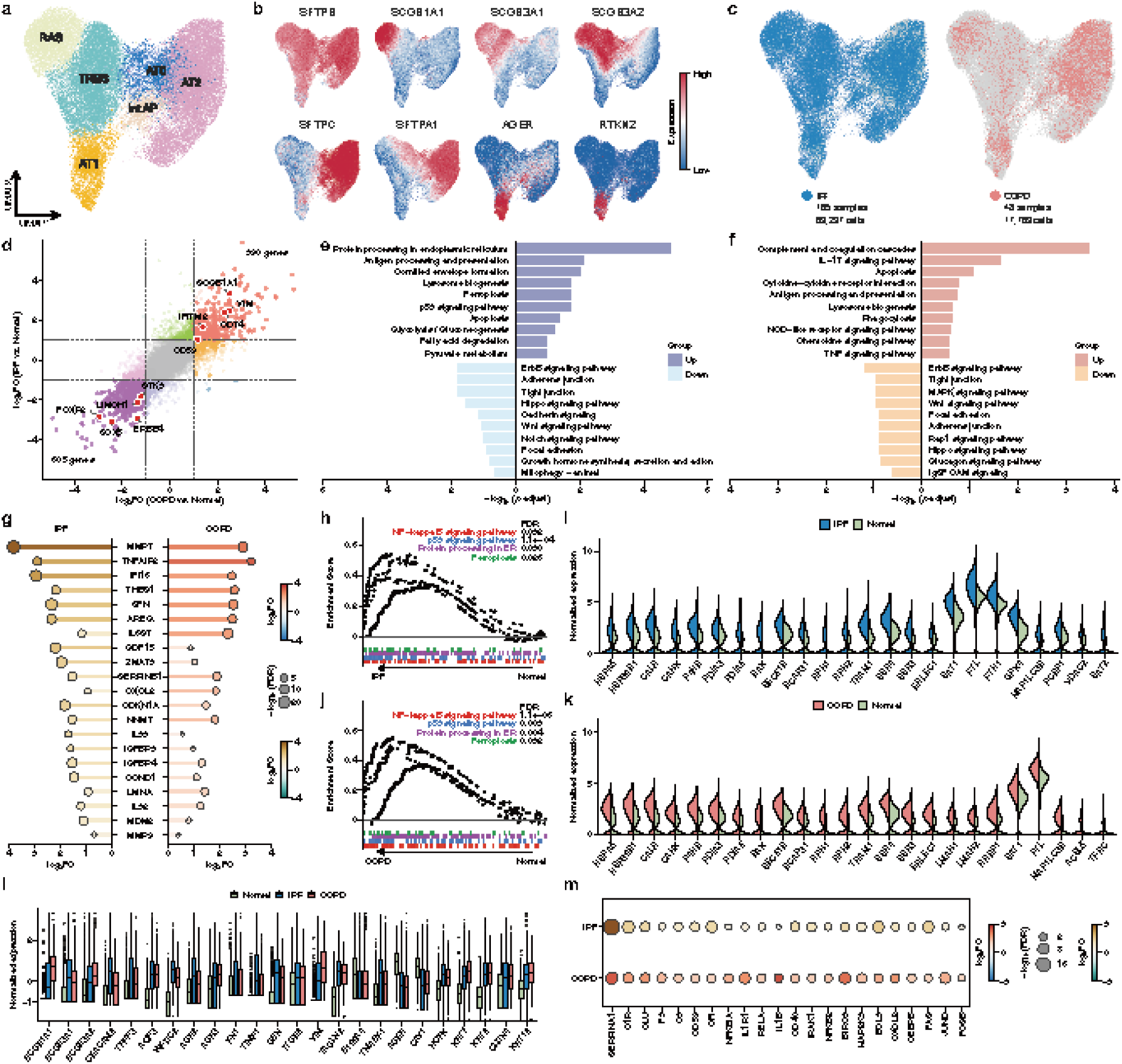
Aberrant Int AP cellular state in human COPD and IPF. **a-c**. UMAP visualization of the diseased lung cell atlas, coloured by cell type annotations (**a**), selected marker gene expression (**b**), and disease status (**c**). **d**. Scatter plot comparing DEGs between normal and diseased Int AP cells. Each point represents one gene. The x axis shows the COPD versus normal comparison, and the y axis shows the IPF versus normal comparison. Statistical significance was assessed using two-sided negative-binomial generalized linear model likelihood-ratio tests, with *P* values adjusted using the Benjamini–Hochberg method. **e-f**. Pathway enrichment analysis of DEGs identified between normal and IPF-derived Int AP cells (**e**) or COPD-derived Int AP cells (**f**). Genes were retained as DEGs only if detected in more than 50% of cells in at least one group. Statistical significance was assessed using a one-sided hypergeometric test, with *P* values adjusted using the Benjamini–Hochberg method. **g.** Fold-change comparison of senescence-associated genes in IPF- and COPD-derived Int AP cells relative to normal controls. Statistical significance was assessed using two-sided negative-binomial generalized linear model likelihood-ratio tests, with *P* values adjusted using the Benjamini–Hochberg method. **h.** GSEA showing enrichment of the NF-κB signaling, p53 signaling, protein processing in ER, and ferroptosis in IPF-derived Int AP cells. Statistical significance was assessed using the fgsea implementation of a two-tailed preranked GSEA, with *P* values adjusted using the Benjamini–Hochberg method. **i.** Split violin plot comparing the expression of representative genes associated with protein processing in ER and ferroptosis. **j.** GSEA showing enrichment of the NF-κB signaling, p53 signaling, protein processing in ER, and ferroptosis in COPD-derived Int AP cells. **k.** Split violin plot comparing the expression of representative genes associated with protein processing in ER and ferroptosis. **l.** Box plot showing the expression of aberrant differentiation-associated genes in Int AP cells derived from normal, IPF and COPD samples. Single-cell expression profiles were used for analysis, with each dot representing a single cell. Center lines indicate medians, box limits indicate the interquartile range, and whiskers indicate 1.5 times the interquartile range. **m.** Dot plot showing fold changes in inflammatory genes in IPF- and COPD-derived Int AP cells relative to normal controls. Statistical significance was assessed using two-sided negative-binomial generalized linear model likelihood-ratio tests, with *P* values adjusted using the Benjamini–Hochberg method.

**Extended data Fig. 8.**
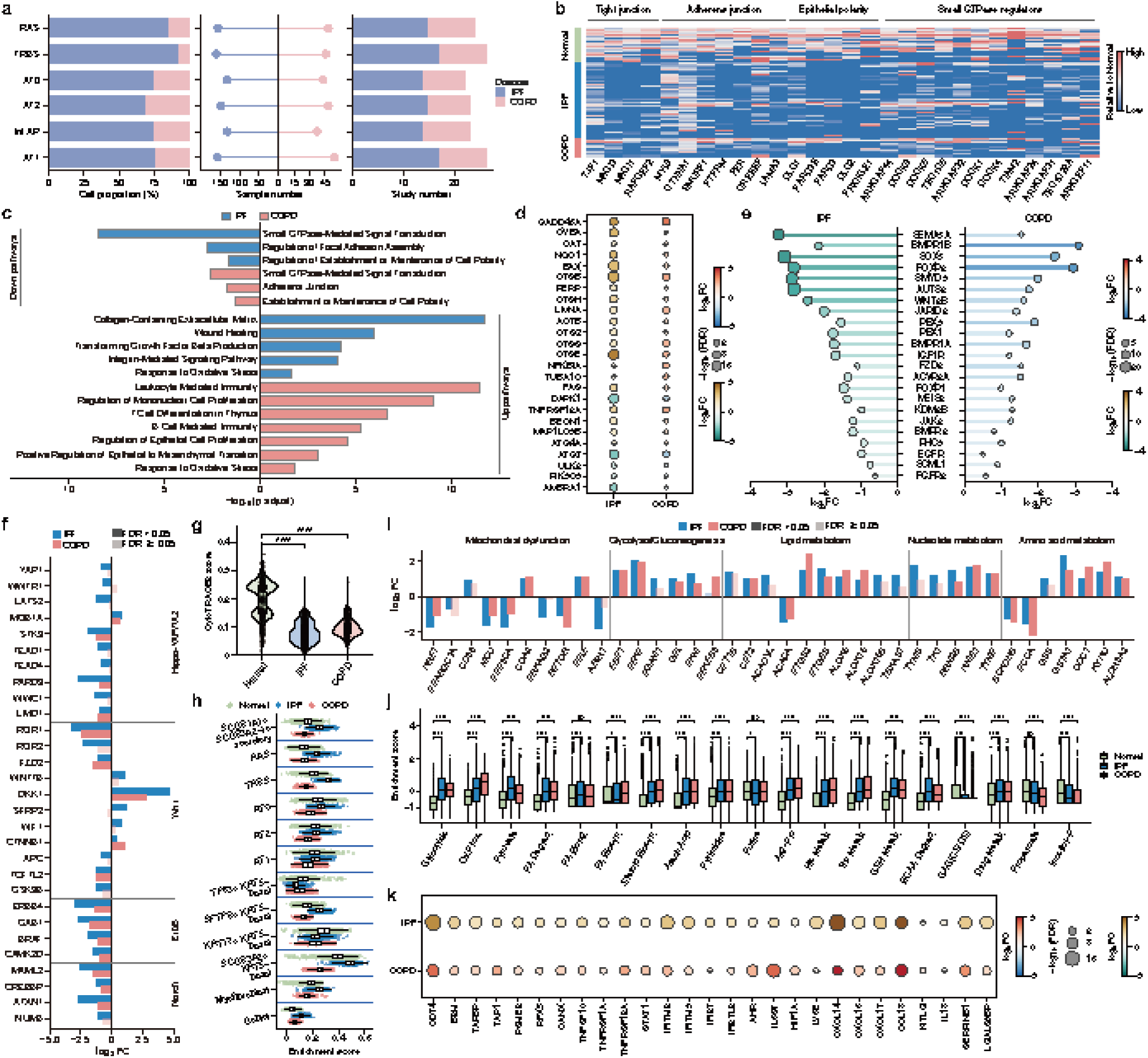
Transcriptional features of dysfunctional Int AP cells. **a.** Number of samples and publications with detected IPF- and COPD-derived Int AP cells. **b.** Heatmap showing the expression of epithelial barrier-associated genes in Int AP cells from normal, IPF and COPD samples. **c.** Bar plot showing GO enrichment analysis of DEGs in IPF- and COPD-derived Int AP cells relative to normal controls. Statistical significance was assessed using a one-sided hypergeometric test, with *P* values adjusted using the Benjamini–Hochberg method. **d.** Dot plot showing fold changes in selected genes in IPF- and COPD-derived Int AP cells relative to normal controls. Statistical significance was assessed using two-sided negative-binomial generalized linear model likelihood-ratio tests, with *P* values adjusted using the Benjamini–Hochberg method. **e.** Fold-change comparison of lung regeneration-associated genes in IPF- and COPD-derived Int AP cells relative to normal controls. Statistical significance was assessed using two-sided negative-binomial generalized linear model likelihood-ratio tests, with *P* values adjusted using the Benjamini–Hochberg method. **f.** Bar plot showing fold changes in genes associated with Hippo-YAP/TAZ, Wnt, ErbB and Notch signalling in IPF- and COPD-derived Int AP cells relative to normal controls. Statistical significance was assessed using two-sided negative-binomial generalized linear model likelihood-ratio tests, with *P* values adjusted using the Benjamini–Hochberg method. **g.** Violin plot showing CytoTRACE2 scores in Int AP cells from normal, IPF and COPD samples. Statistical significance was calculated using a two-sided Wilcoxon rank-sum test. \**P* < 0.05, \*\**P* < 0.01, \*\*\**P* < 0.001, \*\*\*\**P* < 0.0001; ns, not significant. **h.** Direct comparison of gene set score differences in Int AP cells from normal, IPF and COPD lungs. **i.** Bar plot showing fold changes in genes associated with mitochondrial dysfunction and metabolic reprogramming in IPF- and COPD-derived Int AP cells relative to normal controls. **j.** Box plot showing single-cell metabolic activity in Int AP cells from normal, IPF and COPD lungs, as inferred by scMetabolism^74^. Centre lines indicate medians, box limits indicate interquartile ranges and whiskers indicate 1.5 times the interquartile range. Abbreviations: Glycolysis, glycolysis; Glycolysis/Gluconeogenesis, glycolysis and gluconeogenesis; Ox.Phos., oxidative phosphorylation; Pyruvate, pyruvate metabolism; FA Degrad., fatty acid degradation; FA Elong., fatty acid elongation; FA Biosyn., fatty acid biosynthesis; Steroid Biosyn., steroid biosynthesis; Arach.Acid, arachidonic acid metabolism; Pyrimidine, pyrimidine metabolism; Purine, purine metabolism; Arg-Pro, arginine and proline metabolism; His Metab., histidine metabolism; Trp Metab., tryptophan metabolism; GSH Metab., glutathione metabolism; BCAA Degrad., valine, leucine and isoleucine degradation; GAG(CS/DS), glycosaminoglycan biosynthesis (chondroitin sulfate/dermatan sulfate); Drug Metab., drug metabolism–other enzymes; Propanoate, propanoate metabolism; Inositol-P, inositol phosphate metabolism. Statistical significance was calculated using a two-sided Wilcoxon rank-sum test. \**P* < 0.05, \*\**P* < 0.01, \*\*\**P* < 0.001, \*\*\*\**P* < 0.0001; ns, not significant. **k.** Dot plot showing fold changes in inflammatory genes in COPD- and IPF-derived Int AP cells relative to normal controls. Statistical significance was assessed using two-sided negative-binomial generalized linear model likelihood-ratio tests, with *P* values adjusted using the Benjamini–Hochberg method.

Pathway enrichment analysis revealed that Int AP cells were associated with aberrant cell states in both diseases, including cellular senescence, oxidative stress, endoplasmic reticulum stress, autophagy dysfunction, apoptosis, mitochondrial dysfunction, pro-fibrotic matrix remodeling, inhibition of regenerative pathways, epithelial barrier dysfunction and disrupted cell polarity (**Fig. 6e-f** and **Supplementary Table 6**). Compared with normal controls, Int AP cells from COPD and IPF lungs showed downregulation of tight junctions, adherens junctions, IgSF CAM signaling and small GTPase-mediated signal transduction. These changes involved genes encoding the scaffolding protein ZO-1 (*TJP1*), membrane-associated guanylate kinases (*MAGI1* and *MAGI3*) and genes related to polarity establishment (*DOCK4*, *ARHGAP24* and *TIAM2*) (**Extended data Fig. 8b-c**). These transcriptional changes were consistent with impaired epithelial barrier maintenance, cell polarity and cell-cell adhesion. In both COPD and IPF, Int AP cells exhibited activation of senescence-associated p53 and NF-kappaB signalling, together with increased expression of the cell-cycle arrest marker *CDKN1A*, the p53 target gene *SFN*, matrix metalloproteinases (*MMP1*, *MMP7* and *MMP9*) and SASP factors (*GDF15* and *AREG*) (**Fig. 6g**). These results identified a shared senescence-associated inflammatory state in diseased Int AP cells. Additionally, stress-response programmes were also altered in impaired Int AP cells. Growth arrest and DNA damage-inducible alpha (*GADD45A*), the NADPH oxidase complex component *CYBA*, antioxidant defense genes (*CAT* and *NQO1*), pro-apoptotic factors (*BAX*, *CTSB* and *PERP*), and autophagy initiation signals (*BECN1* and *MAP1LC3B*) were upregulated, whereas the autophagy E1 enzyme ATG7 was downregulated (**Extended data Fig. 8d**). Gene set enrichment analysis (GSEA) further showed activation of protein processing in the endoplasmic reticulum (ER) and ferroptosis pathways in Int AP cells from both COPD and IPF. These changes were accompanied by increased expression of unfolded protein response (UPR) and ER protein-related genes (*HSPA5*, *CALR*, *P4HB* and *PDIA3*) and ferroptosis-associated genes (*SAT1* and *FTL*) (**Fig. 6h-k**).

Compared with normal Int AP cells, impaired Int AP cells showed decreased expression of lung developmental TFs (*SOX5*, *FOXP2* and *MEIS2*), epigenetic regulators (*SMYD3* and *AUTS2*), and growth factor receptors (*FGFR2*, *IGF1R*, *BMPR1A* and *BMPR2*), indicating suppression of developmental and regenerative programmes (**Extended data Fig. 8e**). Consistently, lung regenerative pathways, including Wnt, Hippo and ErbB signalling, were enriched among genes downregulated in impaired Int AP cells in both diseases. These genes included *ROR1*, *TCF7L2*, *GSK3B*, *STK3*, *WWC1*, *ERBB4* and *GAB1* (**Extended data Fig. 8f**). In IPF-derived Int AP cells, the Notch signalling transcriptional co-activator *MAML2*, the pathway regulatory switch *CREBBP*, the Hippo pathway effectors *YAP1* and *WWTR1*, and their downstream TFs (*TEAD1* and *TEAD4*) were also downregulated (**Extended data Fig. 8f**). The secretory cell marker *SCGB1A1* was detected in 82.4% of Int AP cells from COPD lungs and 51.5% of Int AP cells from IPF lungs, compared with 1.08% in normal lung (**Fig. 6l**). Other secretory cell markers, including *SCGB3A1*, *AGR2*, *AGR3*, *WFDC2*, *CEACAM6*, *AQP3* and *TPPP3*, were also upregulated in impaired Int AP cells. These findings were consistent with bronchiolization or secretory-like abnormal differentiation. Int AP cells from both COPD and IPF lungs further showed increased expression of fibrotic markers (*COL1A1*, *FN1* and *TIMP1*) and epithelial-mesenchymal transition (EMT)-associated genes (*VIM*, *TAGLN2* and *SFN*). Cornified envelope-associated keratins (*KRT8*, *KRT7* and *KRT18*) were increased in IPF-derived Int AP cells, whereas AT1 differentiation markers (*AGER*, *CAV1* and *HOPX*) were decreased (**Fig. 6l**). Consistent with these transcriptional changes, diseased Int AP cells showed lower CytoTRACE2 scores than normal controls (**Extended data Fig. 8g**). Secretory cell marker gene set scores were increased in diseased Int AP cells, with a stronger shift in IPF than in COPD (**Extended data Fig. 8h**). Together, these results identified impaired regenerative potential and aberrant differentiation programmes in Int AP cells from COPD and IPF lungs.

The expression of the mitochondrial calcium uniporter (*MCU*) and protein phosphatase 3 catalytic subunit alpha (*PPP3CA*) was reduced in diseased Int AP cells, suggesting impaired mitochondrial Ca²⁺ uptake and calcium signalling required for AT2-to-AT1 differentiation (**Extended data Fig. 8i**). Impaired Int AP cells also showed metabolic reprogramming, with increased expression of key enzymes involved in aerobic glycolysis (Warburg effect), including *FBP1*, *PFKP* and *PGAM1*, and carnitine palmitoyltransferases (*CPT1B* and *CPT2*), alongside decreased expression of *ACACA* and *BCKDHB*, rate-limiting enzymes in fatty-acid biosynthesis and branched-chain amino acid catabolism, respectively (**Extended data Fig. 8i**). scMetabolism^74^ analysis further showed enhanced glycolysis, disrupted lipid metabolism and altered amino-acid metabolism in diseased Int AP cells, with stronger metabolic dysregulation in IPF than in COPD (**Extended data Fig. 8j**). Multiple inflammation-related pathways were specifically upregulated in COPD-derived Int AP cells, including complement and coagulation cascades (*C3* and *C1R*), NF-κB signalling (*IL1R1* and *NFKBIA*), TNF signalling (*CXCL2* and *CEBPB*), and the IL-17 signalling pathway (*JUND* and *FOSB*) (**Fig. 6m**). These cells also expressed high levels of inflammatory response-associated molecules, including chemokines (*CXCL14*, *CXCL16* and *CXCL17*), and interferon signalling genes (*STAT1*, *IFITM2* and *IFITM3*) (**Extended data Fig. 8k**). Overall, our data indicate that dysfunctional Int AP cells contribute to aberrant alveolar repair and regenerative failure in both IPF and COPD, and may represent a candidate target for reparative or regenerative cell-based therapies for lung injury and degeneration.

## Discussion

We present here HDLCA, a large-scale single-cell reference atlas of the human respiratory system across the lifespan, by integrating 253 datasets from 225 studies, and comprising more than 18 million cells from 3,198 samples and 2,460 individuals in both health and disease. To our knowledge, it represents the most comprehensive single-cell reference atlas of the human respiratory system assembled to date. This atlas-level integrative resource overcomes the limitations of individual studies, presenting an opportunity to reveal rare, transitional, reparative and disease-associated cell states that may not be captured in the individual datasets. We showcase the identification of Int AP cells as a rare alveolar epithelial progenitor population in normal lungs that shares transcriptional features of both AT2 and AT1 cells, and whose dysfunction may contribute to abnormal alveolar repair and regenerative failure in both IPF and COPD.

Whether the intermediate state co-expressing AT1 and AT2 cell markers observed during AT2-to-AT1 differentiation represents a physiological regenerative intermediate or a consequence of aberrant epithelial repair remains controversial^75–86^. The intermediate alveolar states have been observed in the human IPF lungs and in murine modeling of lung injury, which referred to as alveolar differentiation intermediate (ADI)^37^, pre-alveolar type-1 transitional cell state (PATS)^38^, damage-associated transient progenitors (DATPs)^42^, alveolar-basal intermediates (ABIs)^39^, aberrant basaloid cells^87^, and aberrant intermediate alveolar epithelial cells^40^. These populations are commonly characterized by transitional markers such as *KRT8* and *CLDN4*, EMT-associated genes including *IM*, *FN1* and *COL1A1*, senescence markers including *CDKN1A*, *CDKN2A* and *GDF15*, and basal markers such as *KRT5*, *TP63* and *KRT17*. By contrast, AT1 and AT2 cells arise directly from a bipotent progenitor that shares the transcriptional signature with AT1 and AT2 cells in the murine developing lungs^88,89^, and similar cell populations have been reported in the human fetal lungs^90^.

Contrary to these cells^8,32,36–40,42,88,89^, Int AP cells are detected in both healthy and diseased human lungs, as supported by independent spatial transcriptomic datasets from normal adult lungs^59,60^ and organoid datasets from HLOCA^58^. In normal fetal and adult lungs, Int AP cells co-expressed mature alveolar epithelial markers, including AT2-associated *SFTPC*, *SLC34A2* and *NAPSA* and AT1-associated *AGER*, *CAV1* and *HOPX*, while also sharing transcriptional features marked by alveolar progenitor cell marker *TM4SF1*^91^, growth factor receptors (*TGFBR2* and *FGFR2*) and tip epithelial cell surface markers (*CD44* and *CD36*)^92^. Beyond previously described RAS-to-AT2 cell differentiation^31^, our analysis suggests that both RAS and AT2 cells can potentially give rise to Int AP cells participating in alveolar regeneration. We observed that this intermediate phenotype occurred in normal AT2-to-AT1 differentiation rather than representing a previously reported injury-induced transient state^8,37–39,42^. Int AP cells are associated with genetic susceptibility to both IPF and COPD, and show dysfunction in both diseases. These dysfunctional Int AP cells shared senescence-associated, EMT phenotype and abnormal differentiation programmes with previously described lung injury-induced transient states^93–100^. However, Int AP cells in COPD and IPF lungs expressed secretory cell markers, including *SCGB1A1* and *SCGB3A1*, but lacked basal cell markers such as *KRT5*, *TP63* and *KRT17*. This is similar to the *SCGB3A2*^+^/*SFTPC*^+^ cells observed in human COPD alveoli^12,31^. Kadur *et al*.^59^ revealed that the differentiation of AT2 cells into terminal and respiratory bronchioles secretory (TRBS) cells (*SCGB3A2*^+^/*SFTPB*^+^/*SCGB1A1*^−^/*SFTPC*^−^) or AT1 cells transiently produces *SCGB3A2*^+^/*SFTPC*^+^/*AGER*^+^ cells. Future work will be required to determine whether these observations represent cells arrested in a secretory-to-Int AP transitional state or transdifferentiation toward a secretory fate in impaired alveolar repair.

Recent work has identified AT1 cells as alveolar epithelial mechanosensors^70^, and our findings suggest that Int AP cells may represent an additional mechanosensor in the normal lung. Given that Hippo signalling activity regulates alveolar regeneration^101^ and AT1 cell plasticity^102^, we further found that Hippo-YAP/TAZ signalling governing Int AP lineage specification. Moreover, disease-associated disruption of Hippo signalling and its interacting pathways may contribute to impaired Int AP-mediated alveolar repair. Dysfunction of alveolar epithelial cells, including senescence, defective regeneration, endoplasmic reticulum stress, and apoptosis, constitutes one of the pathological features of IPF and COPD^103–105^. Similarly, impaired epithelial barriers, cellular polarity, and cell-cell junctions in Int AP cells under both diseases lead to a reduced capacity to defend against toxic agents or pathogens. We also found activation of ferroptosis pathways in Int AP cells from both IPF and COPD lungs, which may contribute to aberrant repair of these cells. Inflamed alveolar epithelial cell states have been described in COPD lungs^12,13^, and the stronger inflammatory response of Int AP cells observed in COPD than in IPF. Our observations support the view that Int AP cells represent a rare endogenous alveolar progenitor population whose dysfunction may contribute to the progressive loss of normal alveolar architecture in COPD and IPF. Further lung bioengineering approaches that remove or replace damaged Int AP cells, together with therapeutic strategies that modulate Int AP cell function in injured lungs, may facilitate alveolar regeneration.

However, integrating large-scale single-cell datasets with diverse study designs, experimental conditions, and laboratory sources presents realistic challenges in batch correction^51,106^. We evaluated six existing integration methods, including scANVI^46^ and scPoli^47^, both of which incorporate biological covariates, as well as scVI^45^, Harmony^48^, SCALEX^49^ and Scanorama^50^. Then, scANVI with the best biological variance conservation was selected for constructing the HDLCA. Indeed, both rare and transitional cell populations were identified as distinct clusters, highlighting the high-quality representation provided by HDLCA. In contrast, other methods failed to distinguish these populations, which mix with other clusters due to over-integration. Current batch-correction methods align cell types by modelling relationships across datasets, including batch effects, technical variation and biological differences. Confounded study designs can make it difficult for the model to distinguish biological variation from batch effects^46,107^. When datasets from multiple disease contexts are integrated, disease-specific biological signals may be mistaken for batch effects and consequently attenuated or removed. Future advances in batch-correction methods, balanced experimental designs, or the construction of disease-specific reference atlases will effectively address this limitation.

In summary, we present an atlas-level integrated reference single-cell atlas covering the whole human respiratory system across the lifespan as a living resource to uncover cellular dynamics in development, health and disease, which will facilitate the development of targeted therapeutics strategies for respiratory diseases.

## Methods

### Ethical approval

All public datasets used in this study have published ethics approvals, with detailed information for each study recorded in **Supplementary Table 7**.

### Data collection

We collected publicly available datasets from 271 publications, including human respiratory single-cell transcriptomes, as well as scATAC-seq and spatial transcriptomics datasets (**Supplementary Table 7**). The samples cover the upper respiratory tract (nose, pharynx and larynx), the lower respiratory tract (trachea, bronchi, lungs), organoids, and cell lines. Raw sequencing files (FASTQ, SRA or BAM format), count matrices and processed expression matrices were retrieved from the Gene Expression Omnibus (GEO) (https://www.ncbi.nlm.nih.gov/geo/), the UCSC genome browser (https://genome.ucsc.edu/), ArrayExpress (https://www.ebi.ac.uk/arrayexpress/), CELLxGENE (https://cellxgene.cziscience.com/), and other public repositories. A systematic literature search was performed using the easyPubMed (v2.13)^108^ R package. We searched for potentially relevant studies and datasets by combining respiratory anatomy and disease terms with single-cell and spatial omics terms. Respiratory terms included ‘respiratory’, ‘lung’, ‘trachea’, ‘bronchi’, ‘alveolar’, ‘COVID-19’, ‘SARS’, ‘flu’, ‘COPD’, ‘IPF’, ‘influenza’, ‘smoke’, ‘allergic’ and ‘asthma’. Omics terms included ‘single-cell RNA-seq’, ‘scRNA-seq’, ‘snRNA-seq’, ‘scATAC-seq’ and ‘spatial transcriptomic’. All retrieved records were manually reviewed and curated. Duplicate entries, datasets lacking sufficient metadata, and studies unrelated to the respiratory system were excluded. Only datasets that met the predefined inclusion criteria and were relevant to the scope of this study were retained for downstream analyses.

### Metadata curation and harmonization

Both sample- and cell-level metadata were manually curated and harmonized using standardized naming conventions to ensure consistency across datasets. The harmonized metadata is organized into 21 predefined annotation fields, including sample ID, sample type, sample status, organ, anatomical region, donor gender, donor age, health status, developmental stage, sequencing technology, sequencing method and original cell-type annotation. Standardized metadata were integrated into a unified schema to support dataset indexing, querying and cross-study comparisons. All datasets were archived in independent directories named according to their assigned Dataset IDs. Within each directory, data were organized according to the original accession code, species, omics modality and sequencing platform, providing a structured framework for data storage, management and subsequent analyses.

For sample-level metadata, ‘donor_ID’ and ‘sample_ID’ were assigned as unique identifiers for each donor and sample, respectively. The ‘seq_tech’ and ‘seq_method’ recorded the sequencing technology and sequencing method. The ‘ethnicity’ stored the self-reported ethnicity of each individual. The ‘region’ described both the sample source (such as human tissue, organoid or cell line) and the original sampling location (such as nasal cavity, larynx or lung). Sampling locations were further standardized according to respiratory system anatomical regions and stored in ‘region_normalized’. The ‘donor_gender’ recorded the donor gender as reported in the original study. The ‘donor_status’ represented the health status of the donor, whereas ‘sample_status’ recorded the health status of the sampled anatomical site. Disease annotations followed the standardized nomenclature of Mondo Disease Ontology^109^ and were recorded in ‘mondo_ID’ and ‘mondo_name’. ‘donor_age’ stored donor age, with fetal samples recorded in post-conception weeks (PCW) and postnatal donors recorded in years. For records reported as gestational weeks (GW) in the original publications, we calculated PCW value by subtracting 2 weeks from the reported GW values. ‘sample_type’ described the developmental stage of the sample, including fetal, newborn, child and adult. ‘If_patient’ and ‘treatment’ recorded disease status and treatment information, respectively. The ‘reference’ contains the DOI of the publication from which the sample originated. The ‘project_code’ stored the original dataset accession provided by the source study (such as GSE, SCP or E-MTAB), whereas ‘dataset_id’ served as a unique dataset identifier composed of the publication DOI and a numerical index.

For cell-level metadata, the unique cell barcode and original cell-type annotations were stored in ‘cell_ID’ and in ‘ann_level_1’ to ‘ann_level_5’, respectively, enabling traceability of individual cells to their original annotations.

### Data preprocessing and visualization

We converted raw reads in SRA and BAM formats to FASTQ files using the SRA-toolkit (https://github.com/ncbi/sra-tools) and bamtofastq (https://github.com/10XGenomics/bamtofastq), respectively. FASTQ files were processed using Cellranger (v7.2.0) software with GRCh38 (v3.0.0) genome as a reference (**Supplementary Table 7**). For raw count matrices from Travaglini *et al*.^22^, which were generated using the GRCh38.p12 human reference, we retained only genes overlapping with the GRCh38 (v3.0.0) genome. This filtering retained 20,473 genes from the 10x Genomics profile and 33,233 genes from the Smart-seq2 dataset. For a few processed datasets (normalized matrix), scDenorm (v0.0.9)^110^ was employed to revert them to raw counts. All cell-by-gene count matrices were loaded and processed using the SCANPY (v1.11.4)^111^ Python package. Doublets were predicted for each sample using Scrublet (v0.2.3)^112^ Python package, and cells with doublet score greater than 0.25 were excluded. Cells with fewer than 200 detected genes or more than 20% mitochondrial gene counts were removed, and genes detected in fewer than 3 cells were excluded for downstream analyses. Count matrices were then normalized and log-transformed using the standard SCANPY^111^ workflow.

Before construction of the HDLCA reference atlas, individual studies were analysed separately to generate high-confidence cell type annotations that served as prior labels for scANVI integration. For each study, feature selection, batch correction, dimensionality reduction, clustering and cell-type annotation were performed independently, with analytical strategies tailored to dataset characteristics and sequencing platforms (**Supplementary Table 8**). Highly variable genes (HVGs) were identified using SCANPY^111^ in a batch-aware manner, typically employing the Seurat v3 selection method with ‘sample_ID’ specified as the batch key. Batch effects were corrected using dataset-specific approaches, including Harmony^48^, scVI^45^, and scANVI^46^. Datasets with minimal technical variation were analysed without additional batch correction. Cell populations were identified through Leiden clustering of the corrected latent space and annotated using canonical marker genes, or reference-based label transfer. UMAP embeddings were generated for visualization with the scanpy.tl.umap function in SCANPY^111^.

### Generation of the reference atlas

We curated 13 human normal lung datasets generated using 10x Genomics and Smart-seq2 sequencing platforms from 12 published studies^18,31,36,59,60,113–115^ to build reference atlas (**Supplementary Table 2**). The CellRanger unfiltered matrices ^18–20,31,36,59,60,113,115^ were used as an input for the CellBender (v0.3.0)^116^ algorithm to remove ambient RNA contamination with default parameters. In datasets^17,22,114^ where empty-droplet data were unavailable, ambient RNA correction was performed with scCDC (v1.4)^117^ according to default settings. After individually processed datasets were merged, gene expression counts were normalized using the functions scanpy.pp.normalize_total and scanpy.pp.log1p. Platform-specific normalization parameters were applied, with target_sum set to 1 × 10⁴ for cells generated using 10x Genomics and 1 × 10⁶ for cells generated using Smart-seq2. Feature gene selection was carried out using the DeepTree^118^ to identify highly correlated genes within each dataset. Genes identified as highly correlated in at least two datasets were retained, yielding a final feature set of 3,097 genes.

Data integration of the HDLCA reference atlas was performed using scANVI (v1.0.4)^46^, with raw counts as input. We first pretrained a scVI model on the reference atlas using sample_ID as the batch key. The model used four hidden layers and a 30-dimensional latent representation, and gene expression counts were modelled with a negative binomial distribution. Continuous covariates, including log-transformed total UMI counts (log1p_total_counts) and the percentage of mitochondrial transcripts (pct_counts_mt), were incorporated during training. Covariates were not deeply injected into the network architecture. Layer normalization was applied to both the encoder and decoder, and batch normalization was disabled. All other parameters were kept at their default values. Model training was performed using model.train with a batch size of 740. Validation was performed at each epoch, and early stopping was applied if no improvement was observed for 20 consecutive epochs. The evidence lower bound (ELBO) was used as the performance metric. As scANVI requires prior cell type labels, each dataset in the reference atlas was manually annotated based on unsupervised clustering (**Supplementary Table 8**). scANVI models were then initialized from the trained scVI weights using scvi.model.SCANVI.from_scvi_model, with CellClass provided as the cell type label.

### Consensus auto-annotation with popV

To predict cell type annotations across all datasets, we applied popV (v0.5.3)^34^ using consensus predictions from models pretrained on Tabula Sapiens. The popV framework integrates eight annotation methods, including random forest (RF), support vector machine (SVM), scANVI, OnClass, Celltypist, and k-nearest neighbors implemented following batch correction using three single-cell integration approaches, including scVI, BBKNN and Scanorama. The Tabula Sapiens reference was stratified into four major cell compartments: epithelial, mesenchymal, immune and endothelial cells. For model training, both query_batch_key and ref_batch_key were set to sample_ID, and 300 cells were randomly sampled per cell type label (n_samples_per_label = 300). All other parameters were kept at their default settings.

### Cellular similarity assessment using SCimilarity

Before manual cell-type annotation, SCimilarity (v0.3.0)^35^ was applied to perform cross-dataset searches for human cell similarity. Input single-cell datasets were required to contain a raw count matrix layer, and the officially released pretrained SCimilarity model trained on 23.4 million human cells was used. Cell-type annotation was conducted in the constrained classification mode, in which the reference dataset was subsetted to include only predefined cell types relevant to lung tissue.

The constrained epithelial reference included ‘type I pneumocytes’, ‘type II pneumocytes’, ‘club cells’, ‘goblet cells’, ‘lung ciliated cells’, ‘basal cells’, ‘respiratory basal cells’, ‘secretory cells’, ‘neuroendocrine cells’, ‘pulmonary ionocytes’, ‘squamous epithelial cells’, and ‘mucus-secreting cells’. The mesenchymal reference included ‘pericytes’, ‘vascular-associated smooth muscle cells’, ‘fibroblasts’, ‘myofibroblasts’, ‘smooth muscle cells’, ‘mesothelial cells’, and ‘mesenchymal stem cells’. The endothelial reference included ‘lymphatic endothelial cells’, ‘arterial endothelial cells’, ‘venous endothelial cells’, and ‘capillary endothelial cells’. The immune reference included ‘lung macrophages’, ‘alveolar macrophages’, ‘classical monocytes’, ‘non-classical monocytes’, ‘conventional dendritic cells’, ‘plasmacytoid dendritic cells’, ‘plasmablasts’, ‘mast cells’, ‘CD4⁺ αβ T cells’, ‘regulatory T cells’, ‘CD8⁺ αβ T cells’, ‘mature NKT cells’, ‘natural killer cells’, ‘B cells’, ‘neutrophils’, and ‘plasma cells’. SCimilarity analyses were performed following the official documentation, with default parameters unless otherwise specified.

### Manual cell type annotation

To accurately assign cluster identities in the HDLCA reference atlas, we first identified lineage-specific clusters based on the expression of canonical markers, with *EPCAM*, *PECAM1*, *DCN* and *PTPRC* marking epithelial, endothelial, mesenchymal and immune cell compartments, respectively. Each distinct compartment was then re-clustered to achieve a finer resolution of cell types. Clusters within each HDLCA compartment atlas were annotated using previously reported and cell type-specific marker genes as in **Supplementary Table 1**.

### Benchmarking of data integration methods

To evaluate the performance of HDLCA reference data integration, benchmarking analyses were performed using the GPU-accelerated scib-metrics package (v.0.5.6)^119^. Integration quality was evaluated based on both biological conservation and batch-effect removal. Biological conservation was evaluated using Isolated Labels, normalized mutual information (NMI), Adjusted Rand Index (ARI), Silhouette Label, and cell-type separation (cLISI). Batch correction performance was assessed using Batch average silhouette width (BRAS), batch mixing (iLISI), k-nearest-neighbor batch effect test (KBET), Graph Connectivity, and Principal Component Regression (PCR) comparison. The performance of Harmony^48^, SCALEX^49^, scVI^45^, scANVI^46^, and scPoli^47^, Scanorama^50^ was compared using a common batch key (‘sample_ID’). Method-specific configurations included: SCALEX (min_cells = 1, min_features = 1), scVI (default parameters, 10 epochs), scANVI (label_key = ‘celltype’, 10 epochs), scPoli (unsupervised mode, 10 epochs), and Scanorama (dimred = 10). All other parameters were set to their default values.

### Accuracy evaluation of cell clustering

Clustering robustness and cell-type annotation accuracy of the HDLCA reference atlas were evaluated using SCCAF^33^ method. The scANVI latent embedding was used as the feature matrix, with cell-type annotations serving as class labels. The support vector machines (SVM) classifier was trained using up to 500 randomly selected cells per cell type, with all other parameters kept at their default settings.

### Extension of reference atlas by transfer learning

The extended datasets were projected onto the reference atlas using the transfer-learning method scArches^44^ as implemented in scvi-tools (v1.0.4)^120^. Cell-type annotations from the HDLCA reference atlas were used to guide label transfer and integration of newly collected datasets into the shared latent space. The model was trained with four hidden layers using 128 nodes and 30 latent dimensions, using ‘sample_ID’ as the batch covariate and HDLCA annotated cell identities as the reference label. Log-transformed total UMI counts (log1p_total_counts) and the percentage of mitochondrial transcripts (pct_counts_mt) were included as continuous covariates. All remaining parameters were set to their default values.

### Construction of the diseased cell atlas

To construct the diseased cell atlas, a subset of cells annotated as distal epithelial cells derived from patients with COPD and IPF in the HDLCA was selected. Following gene intersection across all datasets, 11,870 genes were retained for downstream analysis. Studies contributing fewer than 50 cells were excluded prior to feature selection. Top 500 HVGs were selected per study in the ‘seurat_v3’ flavor, with a loess smoothing span of 0.8. Principal component analysis (PCA) was performed on the HVG-subset expression matrix. Batch-effect correction was then applied using Harmony^48^ approach, with individual studies as batch key, yielding a corrected low-dimensional embedding. The neighborhood graph of cells was subsequently computed in the Harmony-corrected PCA space using 10 principal components. For visualization, dimensionality reduction was performed using UMAP with the scanpy.tl.umap function with default parameters. Unsupervised clustering using the Leiden algorithm revealed six cell types (RAS, TRBS, AT0, AT2, Int AP and AT1) that were reannotated based on the expression of canonical marker genes (**Supplementary Table 1**).

### Pseudobulk profiles generation

For HDLCA reference atlas, pseudobulk gene expression profiles were generated from raw counts stored in the ‘counts’ layer of the AnnData object using scanpy.get.aggregate in SCANPY (v1.11.5)^111^. Cells were grouped by HDLCA cell type annotation (‘CellType_HDLCA’) and sample identifier (‘sample_ID’). For each gene, raw counts from all cells assigned to the same cell type within each sample were summed to generate pseudobulk count matrices for downstream analyses.

For epithelial and disease-related analyses, pseudobulk matrices were generated using the AggregateExpression function in Seurat (v5.2.1)^121^. In short, cells were grouped by sample identity together with the corresponding cell-state, disease or source label, and raw counts for each gene were summed within each group. For differential analyses of Int AP cells derived from normal, IPF, and COPD samples, SRR14851096 was excluded prior to aggregation due to low quality. The resulting raw pseudobulk count matrices were exported for downstream differential expression analysis.

### Differential gene expression analysis

Differential gene expression analysis was performed on pseudobulk count matrices using edgeR (v4.0.1)^122^. Different groups were specified for differential analysis under various conditions. In each comparison, genes were first filtered according to their expression proportion in the raw single-cell data. The expression proportion was defined as the percentage of cells with RNA counts greater than zero within a given comparison group. Genes were retained if they exceeded the specified threshold in at least one comparison group (min_pct = 0 or min_pct = 0.5). The filtered pseudobulk count matrices were converted into DGEList objects and normalized using the trimmed mean of M-values (TMM) method via calcNormFactors. Dispersion was estimated with estimateDisp, and negative-binomial generalized linear models were fitted using glmFit. Differential expression was tested using likelihood-ratio tests with glmLRT. To account for sample-level effects, sample ID was included as a covariate when fitting the GLM where applicable. For comparisons between disease groups, sample ID was not included because it was collinear with disease status. Genes with a false discovery rate (FDR) < 0.05 and an absolute log2 fold change > 1 were retained for visualization and downstream enrichment analyses.

In the section of association of Int AP with pulmonary disease risk, we performed differential expression analysis of HDLCA-annotated cell populations using the rank_genes_groups function in SCANPY with the Wilcoxon rank-sum test. We defined DEGs as genes with FDR < 0.05 and |log2 fold-change| > 1.

### Gene regulatory network analysis

Gene regulatory network (GRN) analysis was performed using the pySCENIC workflow implemented in OmicVerse (v1.7.7)^123^. Raw count matrices from the epithelial, mesenchymal, endothelial and immune cell compartments were used as input, with genes exhibiting zero expression variance across all cells excluded prior to analysis. A raw co-expression-based GRN was constructed using the scenic_obj.cal_grn function in RegDiffusion under default parameter settings. We downloaded human cisTarget ranking databases (mc_v10_clust) and TF-motif annotation file (motifs-v10nr_clust-nr.hgnc-m0.001-o0.0.tbl) for the next regulon pruning step. Direct TF-target relationships were subsequently inferred using scenic_obj.cal_regulons, with dropout handling enabled during correlation calculation. Regulon activity in each cell was quantified using AUCell. To identify cell-type-specific regulons, we calculated regulon specificity scores (RSS) from the regulon activity matrix and used Jensen-Shannon divergence to compare regulon activity distributions across cell types. We then used the ov.single.cosg function to identify marker regulons for each cell type.

### RNA velocity analysis

RNA velocity analysis was conducted by using velocyto (v0.17.17)^124^ and scVelo (v0.2.5)^62^ Python package. Briefly, the velocyto pipeline was applied to quantify spliced and unspliced transcripts for each sample, and loom files were generated using the ‘run10x’ option. The resulting loom files were merged with the corresponding scRNA-seq AnnData objects using the ‘scv.utils.merge’ function in scVelo. Based on the velocities estimated by the ‘scv.tl.velocity’ function, the velocity graph of cell-to-cell transition probabilities was constructed by the ‘scv.tl.velocity_graph’ function. RNA velocities were estimated with ‘scv.tl.velocity’, followed by construction of the cell-to-cell transition probability graph using ‘scv.tl.velocity_graph’. Finally, the velocity graph was projected onto the UMAP embedding and visualized as streamlines on a regular grid with ‘scv.pl.velocity_embedding_stream’.

### Trajectory inference

Distal epithelial cell differentiation trajectories were reconstructed using Diffusion Map^65^, PAGA^64^, Slingshot^66^, Palantir^67^, SLICE^68^, and Monocle3^63^. Diffusion map embeddings were generated in SCANPY^111^ using the integrated epithelial atlas latent representation and a neighborhood graph constructed with 20 nearest neighbors. To infer connectivity among epithelial cell states, PAGA analysis was performed using the scanpy.tl.paga function with default parameters. The same neighborhood graph used for diffusion map construction was retained for PAGA analysis. Graph visualization and edge filtering were generated using pl.paga_compare with threshold = 0.005 and node_size_scale = 1. Slingshot (v2.18)^66^ trajectory inference was performed using the first two diffusion components and cellular identities without predefined root or terminal states. Palantir^67^ trajectories were inferred using OmicVerse (v1.7.7)^123^, with RAS cells specified as the initial state and AT1 or AT2 cells as terminal states. Cellular differentiation potential was further assessed by calculating scEntropy scores using the bootstrap implementation of SLICE.

### Developmental potential assessment

The differentiation states of distal epithelial cells were inferred by CytoTRACE2^72^ Python package. Gene expression matrices and corresponding cell-type annotations were exported from the integrated distal epithelial cell atlas and provided as input to the CytoTRACE2^72^ framework. Analyses were performed using the full model with a batch size of 10,000 cells and a smoothing batch size of 1,000 cells. Parallelization was disabled, and a maximum of 200 principal components was used for dimensionality reduction. A random seed of 14 was specified to ensure reproducibility. All remaining parameters were retained at their default settings.

### Trajectory-based differential expression analysis

To reduce computational burden while preserving cellular composition, cells were randomly downsampled to 10% of the original dataset in a cell type-stratified manner. Trajectory-dependent differential expression analysis was subsequently performed using tradeSeq (v1.24)^69^ R package. Following downsampling, lineage inference was reconstructed using Slingshot (v2.18)^66^, and the resulting lineage assignments together with the raw count matrix were used as inputs for downstream analyses. The optimal number of knots for generalized additive model (GAM) fitting was determined using the evaluateK function. Pseudotime values and lineage assignment weights for each cell were extracted using slingPseudotime (na = FALSE) and slingCurveWeights, respectively. Gene expression dynamics along trajectories were modeled using fitGAM, incorporating sample_ID as a covariate to account for sample-specific effects and using seven knots. Differential expression patterns between lineages were identified using the patternTest function with default parameters. Genes showing significant lineage-dependent expression dynamics were further grouped into co-expression modules using clusterExpressionPatterns, with nPoints = 40 to capture temporal expression patterns along pseudotime.

### Gene signature scoring

The functional gene signature sets were collected from databases including GO, KEGG and Molecular Signature Database (MsigDB)^125,126^. Gene sets associated with airway cell type (*SCGB1A1*^+^ *SCGB3A2*-lo secretory, RAS, TRBS, AT0, AT2, AT1, *TP63*^+^/*KRT5*^−^ basal, *SFTPB*^+^/*KRT5*^−^ basal, *KRT17*^+^/*KRT5*^−^ basal and *SCGB3A2*^+^/*KRT5*^−^ basal), mucus-secreting cell types (goblet and mucous) and myofibroblast from the HDLCA reference atlas were defined using marker genes (FDR < 0.05 and |log2 fold-change| > 1) identified by the scanpy.pl.rank_genes_groups function with the Wilcoxon rank-sum test. Gene set scores were calculated for each cell using the scanpy.tl.score_genes function.

### Functional enrichment analysis

For the DEGs of each group, GO and KEGG enrichment analyses were performed separately for up- and down-regulated gene sets using the enrichGO_run and enrichKEGG_run wrapper functions from the RNAseqStat (https://github.com/xiayh17/RNAseqStat) R package, with org.Hs.eg.db as the human gene annotation reference. GSEA was performed using enrichgesKEGG_run from the RNAseqStat package, with the full ranked gene list ordered by log2 fold change as input. For genes with different expression patterns between lineages, GO and KEGG enrichment analyses were conducted using the R package clusterProfiler^127^.

### Quantitation of metabolic activity

Single-cell metabolic pathway activities were quantified using scMetabolism^74^ algorithm. Pathway activity scores for individual cells were estimated using the AUCell (Area Under the Curve) method, which evaluates the enrichment of pathway-specific gene signatures at the single-cell level. KEGG was selected as the metabolic type, and all other parameters were set to their default values.

### The development of sc2HDLCA toolkit

The sc2HDLCA toolkit is a computational framework developed to facilitate the use of the integrated HDLCA reference atlas. It enables projection of new datasets by query-to-reference mapping, de novo cell-type annotation, cross-modal integration and deconvolution of spatial and bulk data, with the HDLCA reference atlas serving as the reference for all analyses. The toolkit implements scArches (v0.6.1)^44^ to project new single-cell datasets onto the integrated HDLCA reference atlas. For cell-type annotation, Cell Marker Accordion^128^ was integrated to enable marker-based annotation using HDLCA-defined cell-type markers, and the popV^34^ pipeline was incorporated to support automated annotation based on HDLCA reference atlas. With the integrated atlas as references, sc2HDLCA integrates SCCAF-D^129^ for bulk data deconvolution, as well as RCTD (v2.2.1)^130^ and SpatialDWLS (v1.1.2)^131^ for spatial transcriptomic deconvolution. In addition, the toolkit incorporates uniPort (v1.3)^132^ for the integration of HDLCA reference data with scATAC-seq profiles. All methods were run using default parameter settings.

### Visium data processing and analysis

We downloaded the publicly available Visium spatial transcriptomic datasets of fetal lungs at 12-20 post-conception weeks (pcw) from He *et al*.^20^ and Sountoulidis et al.^17^, as well as adult lungs from Madissoon *et al*.^60^. These spatial transcriptomic datasets include proximal airways and distal lungs, such as trachea, bronchi, low left parenchyma, and top left parenchyma. The processed data were subsequently standardized and organized into AnnData objects using SCANPY (v1.11.4) ^111^ for downstream analyses.

Cell-type abundances in spatial transcriptomic data were inferred using cell2location (v0.1.4)^133^, which integrates single-cell reference transcriptomes with spatial gene expression profiles through a hierarchical Bayesian model. Cell-type specific reference signatures were estimated from the HDLCA reference atlas using annotated cell identities while accounting for sample-level batch effects. The trained model was subsequently used to deconvolve each spatial location and estimate cell-type abundance. Reference and spatial models were trained for 100 and 5,000 epochs, respectively, with an expected 30 cells per location. Unless otherwise specified, default parameters were used. Cell-type abundance maps were generated from the inferred absolute mRNA contributions, and robust abundance estimates were defined as the 5% posterior quantile of the cell-type specific mRNA-count distribution.

For cell type co-localization analysis, we followed the standard pipeline of the mistyR^61^ package to perform cell-type colocalization analysis on Visium spatial transcriptomic datasets with cell types deconvoluted by cell2location^133^.

### Collection and analysis of GWAS summary statistics

We followed the standard pipeline of gsMap (v1.73.4)^53^ to integrate ST data with GWAS summary statistics to map cells to human pulmonary diseases. The GWAS summary statistics were collected from GWAS Catalog (https://www.ebi.ac.uk/gwas/), and converted into the gsMap input format. All GWAS data used in this study are summarized in **Supplementary Table 3**.

To assess disease-associated genetic risk at single-cell resolution, scDRS^57^ was applied to the distal epithelial cell atlas using COPD- and IPF-associated GWAS summary statistics. For each cell, scDRS computed disease relevance scores and corresponding p-value reflecting enrichment of disease-associated genetic signals. Technical and sample-level covariates, including the number of detected genes and sample identity, were incorporated into the model. Because the expression matrices had already been preprocessed, we set ‘flag_filter_data’ to ‘FALSE’ and ‘flag_raw_count’ to ‘FALSE’ to disable additional filtering and raw-count normalization.

### Web portal development

The HDLCA web portal was developed as a static website using Jekyll (v4.3.3; https://jekyllrb.com/) and hosted on GitHub Pages (https://pages.github.com/). The external Data Viewer was developed as a single-page application using Vue.js (v2.7.16; https://v2.vuejs.org/) and deployed on an Nginx (v1.18.0; https://nginx.org/) server running Ubuntu 22.04.2 LTS (https://ubuntu.com/). Interactive tables and data visualization modules were implemented using DataTables (v1.10.21 and v1.12.1; https://datatables.net/), D3.js (v7.9.0; https://d3js.org/), ECharts (v5.4.3; https://echarts.apache.org/) and Plotly.js (v3.7.0; https://plotly.com/javascript/), whereas WebGL-based single-cell scatterplots in the Data Viewer were rendered using regl (v2.1.1; https://github.com/regl-project/regl).

## Data availability

Data included in the HDLCA can be downloaded at https://dlca.gznl.org/ without login. All publicly available datasets analysed in this study were obtained from their original repositories using the accession numbers reported in the corresponding publications. A complete list of datasets, accession numbers, and source repositories is provided in **Supplementary Table 7**.

## Code availability

Code for reproducing this study is available at https://github.com/rnacentre/HDLCA-reproducibility.

## Acknowledgements

We thank Beijing Cloudna Technology Co., Ltd for technical support. We thank Lahong Xu, Yin Huang, Xiaofeng Chen, Haoyu Li, Yangyi Ren and Baowei Huang for data collection. This work is supported by the Major Project of Guangzhou Laboratory (Grant No. GZNL2024A01002, GZNL2023A01006, GZNL2025C01007, HWYQ23-003, YW-YFYJ0102), the National Key R&D Programs of China (2025YFE0200600, 2023YFF1204700, 2024YFF1206600), the National Natural Science Foundation of China (32270707) to Z.M., and the Young Scientists Fund of the National Natural Science Foundation of China (32500586) to L.H., and the Postdoctoral Research Project Funding of Guangzhou (BSHF23-094) to J.W.

## Contributions

L.H. conceived and led the study. K.F., L.J.H., L.Y.H., Z.H., S.F., L.H., J.W., Y.A. and H.S. collected and curated data. J.W. and S.F. built the web portal. L.H. wrote the manuscript. Z.H., S.F., L.H., J.W., K.F., Y.A. and J.Z. performed analyses. Z.M. supervised the work.

## Ethics declarations

### Competing interests

The authors declare no competing interests.

## Notes

### Competing Interest Statement

The authors have declared no competing interest.

### Summary of Updates

Funding information has been updated; author affiliations updated; Parts of the main text and the Methods section have been updated.

